# Dynactin binding to tyrosinated microtubules promotes centrosome centration in *C. elegans* by enhancing dynein-mediated organelle transport

**DOI:** 10.1101/130104

**Authors:** Daniel José Barbosa, Joana Duro, Dhanya K. Cheerambathur, Bram Prevo, Ana Xavier Carvalho, Reto Gassmann

## Abstract

The microtubule-based motor dynein generates pulling forces for centrosome centration and mitotic spindle positioning in animal cells. How the essential dynein activator dynactin regulates these functions of the motor is incompletely understood. Here, we dissect the role of dynactin’s microtubule binding activity, located in p150’s CAP-Gly domain and an adjacent basic patch, in the *C. elegans* zygote. Using precise mutants engineered by genome editing, we show that microtubule tip tracking of dynein-dynactin is dispensable for targeting the motor to the cell cortex and for generating cortical pulling forces. Instead, p150 CAP-Gly mutants inhibit cytoplasmic pulling forces responsible for centration of centrosomes and attached pronuclei. The centration defects are mimicked by mutations of the C-terminal tyrosine of α-tubulin, and both p150 CAP-Gly and tubulin tyrosination mutants decrease the frequency of organelle transport from the cell periphery towards centrosomes during centration. In light of recent work on dynein-dynactin motility *in vitro*, our results suggest that p150 GAP-Gly domain binding to tyrosinated microtubules promotes initiation of dynein-mediated organelle transport in the dividing embryo, and that this function of dynactin is important for generating robust cytoplasmic pulling forces for centrosome centration.

## INTRODUCTION

Cytoplasmic dynein 1 (dynein) is the major microtubule (MT) minus-end directed motor in animals and transports various cargo from the cell periphery to the cell interior. The motor also moves and positions intracellular structures such as nuclei and centrosomes by pulling on the MTs to which they are attached. To generate pulling force, dynein is either attached to anchor proteins fixed at the cell cortex (cortical pulling) (Nguyen-Ngoc *et al*, 2007; Kotak & Gönczy, 2013), or dynein is anchored on organelles in the cytoplasm (cytoplasmic pulling) (Tanimoto *et al*, 2016; Kimura & Kimura, 2011; Wühr *et al*, 2010; Kimura & Onami, 2005; Shinar *et al*, 2011). In the latter instance, dynein generates length-dependent pulling forces by working against viscous drag as it moves organelles along MTs toward centrosomes.

Dynactin is an essential multi-subunit activator of dynein that forms a tripartite complex with the motor and cargo-specific adaptors proteins (Schroer & Sheetz, 1991; Gill *et al*, 1991; Splinter *et al*, 2012; McKenney *et al*, 2014; Schlager *et al*, 2014), but how dynactin supports the diverse functions of dynein remains incompletely understood. Dynactin is built around a short actin-like Arp1 filament and has its own MT binding activity, which resides at the end of a long projection formed by the largest subunit p150 (Schroer, 2004). p150 has a tandem arrangement of MT binding regions, consisting of an N-terminal Cytoskeletal Associated Protein Glycine-rich (CAP-Gly) domain and an adjacent patch rich in basic residues (Waterman-Storer *et al*, 1995; Culver-Hanlon *et al*, 2006). The CAP-Gly domain binds to MTs and to the MT plus-end tracking proteins (+TIPs) CLIP-170 and end-binding (EB) protein. In animal cells, +TIP binding of dynactin recruits dynein to growing MT ends (Watson & Stephens, 2006; Duellberg *et al*, 2014; Vaughan *et al*, 2002; 1999; Splinter *et al*, 2012).

The p150’s CAP-Gly domain recognizes the C-terminal EEY/F motif present in α-tubulin and EB/CLIP-170 (Hayashi *et al*, 2005; Honnappa *et al*, 2006; Weisbrich *et al*, 2007; Steinmetz & Akhmanova, 2008). The C-terminal tyrosine of α-tubulin can be removed and re-ligated in a tyrosination-detyrosination cycle and is proposed to regulate the interactions with molecular motors and other MT binding proteins (Yu *et al*, 2015; Janke, 2014). Tubulin tyrosination is required in mouse fibroblast to localize CAP-Gly proteins, including p150, to MT plus ends (Peris *et al*, 2006), and recent work *in vitro* demonstrated that the interaction between p150’s CAP-Gly domain and tyrosinated MTs enhances the initiation of processive dynein motility (McKenney *et al*, 2016),

The functional significance of MT binding by p150 is best understood in neurons. Single point mutations in the CAP-Gly domain cause the ALS-like motor neuron degenerative disease HMN7B and a form of parkinsonism known as Perry syndrome (Puls *et al*, 2003; Farrer *et al*, 2009; Araki *et al*, 2014). Cellular and *in vivo* studies addressing the underlying molecular defects revealed that p150 CAP-Gly domain-dependent binding of dynactin to dynamic MTs in the distal axon enhances the recruitment of dynein, which in turn facilitates efficient initiation of retrograde transport (Lloyd *et al*, 2012; Moughamian & Holzbaur, 2012; Moughamian *et al*, 2013).

While the critical role of p150’s CAP-Gly domain in neuronal trafficking is firmly established, little is known about how MT binding by dynactin regulates dynein functions in other cellular contexts. A study in *Drosophila* S2 cells observed multipolar spindles with a *glued* construct lacking the CAP-Gly domain, suggesting a role in organizing MT arrays (Kim *et al*, 2007). In budding yeast, introduction of the motor neuron disease mutation into the p150 homolog Nip100 produced a specific defect in the initial movement of the spindle and nucleus into the bud neck during mitosis (Moore *et al*, 2009), suggesting that dynactin binding to MTs helps dynein generate pulling forces under load. In budding and fission yeast, dynein is off-loaded to cortical anchors via MTs for subsequent force production, and in budding yeast this requires MT plus end tracking of dynein. Whether dynein-dependent pulling forces are mechanistically coupled to MT tip tracking of the motor in animals remains to be determined.

MT binding of dynactin is significantly enhanced by electrostatic interactions between the p150 basic patch and the acidic tails of tubulins (Wang *et al*, 2014; Mishima *et al*, 2007; Culver-Hanlon *et al*, 2006). In the filamentous fungus *A. nidulans*, deletion of the basic patch diminishes the accumulation of dynactin and dynein at MT tips and partially impairs nuclear migration and early endosome distribution (Yao *et al*, 2012). Interestingly, humans express tissue-specific splice isoforms of p150 that lack the basic patch (Dixit *et al*, 2008; Zhapparova *et al*, 2009), but the implications for dynactin function are unclear.

In the *C. elegans* one-cell embryo, dynein and dynactin are essential for centrosome separation, migration of the maternal and paternal pronucleus, centration and rotation of the nucleus-centrosome complex, assembly and asymmetric positioning of the mitotic spindle, chromosome congression, and transversal spindle oscillations in anaphase (Gönczy *et al*, 1999; Rose & Gönczy, 2014; Schmidt *et al*, 2005). Here, we use a set of p150^DNC-1^ and α-tubulin mutants constructed by genome editing to define role of dynactin’s MT binding activity in this system. Our results uncover a functional link between the efficient initiation of dynein-mediated organelle transport, which requires dynactin binding to tyrosinated MTs, and the cytoplasmic pulling forces responsible for centration of centrosomes.

## RESULTS

### Identification of +TIPs required for microtubule plus-end targeting of dynein-dynactin in the *C. elegans* early embryo

To investigate whether dynactin’s MT binding activity contributes to dynein function in the early *C. elegans* embryo, we first asked whether dynactin is present at MT plus ends at this developmental stage. Live confocal imaging in the central plane of metaphase one-cell embryos co-expressing endogenous GFP::p50^DNC-2^ and transgene-encoded EBP-2::mKate2 revealed that dynactin travelled on growing MT tips from mitotic spindle poles to the cell cortex (Fig. 1A; Movie S1). Imaging of the cortical plane allowed end-on visualization of MT tips as they arrived at the cortex (Fig. 1B), which facilitated quantification of dynactin levels at plus ends. Measurements of fluorescence intensity revealed the expected positive correlation between GFP::p50^DNC-2^ and EBP-2::mKate2 levels, but also showed that there is considerable variation in the amount of GFP::p50^DNC-2^ at MT plus ends (Fig. S1A). Cortical residency times for EBP-2::mKate2 and GFP::p50^DNC-2^ were nearly identical (1.67 ± 0.03 s and 1.50 ± 0.05 s, respectively) and agreed with previously published measurements for cortical residency times of GFP::EBP-2 (Fig. S1C, D) (Kozlowski *et al*, 2007). We also generated a *dynein heavy chain*^*dhc-1*^*::gfp* knock-in allele to assess the localization of endogenous dynein. DHC-1::GFP was readily detectable on growing MT plus ends in early embryos (Fig. S1F; S4; Movie S2), although the signal was weaker than that obtained for GFP::p50^DNC-2^. We conclude that a pool of dynein-dynactin tracks with growing MT plus ends in the early *C. elegans* embryo.

**Figure 1:**
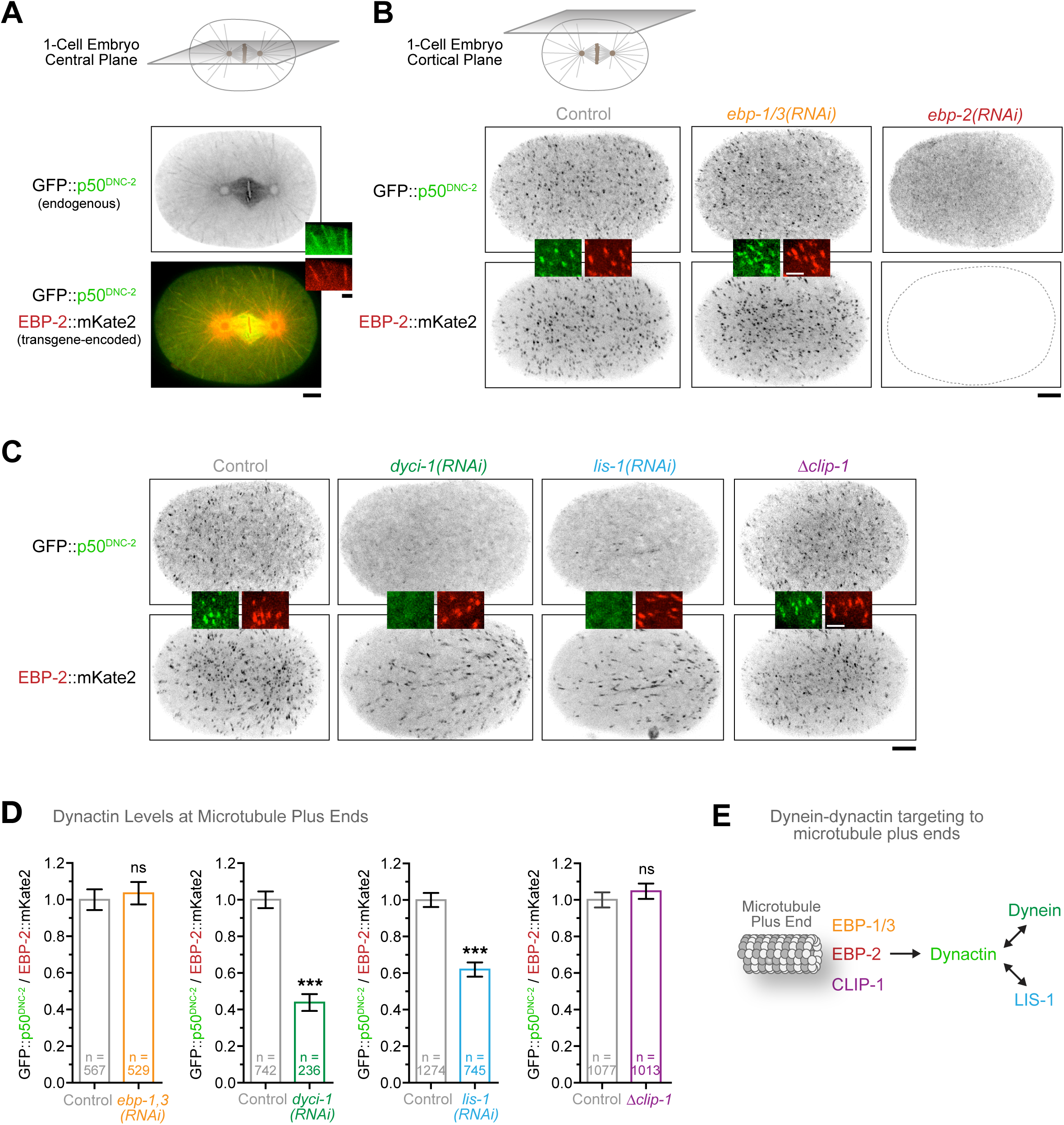
Identification of +TIPs required for MT plus-end targeting of dynein-dynactin in the *C. elegans* early embryo. **(A)** Central confocal section in a *C. elegans* one-cell embryo at metaphase co-expressing GFP::p50^DNC-2^ and EBP-2::mKate2, showing dynactin enrichment at growing MT plus-ends, kinetochores, and the spindle. Image corresponds to a maximum intensity projection over time of 10 images acquired every 200 ms. Scale bar, 5 *μ*m; insets, 2 *μ*m. **(B)** Cortical confocal section in one-cell embryos as in *(A)* without treatment (control) and after *ebp-1/3(RNAi)* or *ebp-2(RNAi)*. Image corresponds to a maximum intensity projection over time of 12 images acquired every 5 s. Scale bar, 5 *μ*m; insets, 2 *μ*m. **(C)** Embryos as in *(B)* without treatment, after *dyci-1*(RNAi) or *lis-1*(RNAi), and with a *clip-1* null allele (Δ*clip-1*). Scale bar, 5 *μ*m; insets, 2 *μ*m. **(D)** Quantification of dynactin levels at MT plus ends using fluorescence intensity measurements of GFP::p50^DNC-2^ at the cortex in the conditions shown in *(B)* and *(C)*. For each MT end, GFP::p50^DNC-2^ signal was normalized to EBP-2::mKate2 signal. Error bars represent the SEM with a 95 % confidence interval. For each condition, *n* indicates the total number of individual MT ends measured in 7-11 embryos. The t-test was used to determine statistical significance (*** indicates p < 0.0001; ns = not significant, p > 0.05). **(E)** Cartoon depicting the MT plus-end recruitment pathway for dynein-dynactin in the early *C. elegans* embryo, based on the results in *(B)*-*(D)*.

We used the quantitative cortical imaging assay to determine which +TIPs were required for MT plus-end targeting of dynactin and dynein. RNAi-mediated depletion of the three EB paralogs revealed that EBP-2 is required for GFP::p50^DNC-2^ targeting to MT plus ends, while EBP-1 and EBP-3 are dispensable (Fig. 1B, D). In mammalian cells, CLIP-170 acts as an essential linker between EB and dynactin (Lansbergen *et al*, 2004; Bieling *et al*, 2008; Akhmanova & Steinmetz, 2015). To assess whether the CLIP-170-like protein CLIP-1 recruits dynactin to MT plus ends in *C. elegans*, we generated a null allele of *clip-1* in the *gfp::p50*^*dnc-2*^ background (Fig. S1E). This revealed that CLIP-1 is dispensable for MT plus-end localization of GFP::p50^DNC-2^ (Fig. 1C, D), suggesting that dynactin is directly recruited by EBP-2. Next, we depleted dynein intermediate chain^DYCI-1^ and the dynein co-factor LIS-1. In both cases, GFP::p50^DNC-2^ levels at MT plus ends decreased substantially (Fig. 1C, D; note that we performed partial depletions because penetrant depletions resulted in sterility of the mother). Conversely, depletion of p150^DNC-1^ showed that DHC-1::GFP targeting to MT plus ends was dependent on dynactin (Fig.S1F, G).

We conclude that in the *C. elegans* early embryo, dynein and dynactin are interdependent for targeting to MT plus ends and require EBP-2 and LIS-1, but not EBP-1, EBP-3, or the CLIP-170 homolog CLIP-1 (Fig. 1E).

### Splice isoforms of the p150^DNC-1^ basic patch modulate microtubule plus-end targeting of dynactin

Having established that dynein and dynactin require the EB homolog EBP-2 for targeting to MT tips, we next examined the role of the dynactin subunit p150^DNC-1^, whose N-terminal CAP-Gly domain (residues 1-69) mediates binding to EB and MTs (Fig. 2A). In addition, p150^DNC-1^ features a ∼200-residue basic-serine rich region between the CAP-Gly domain and the first coiled-coil region, which has been proposed to regulate p150^DNC-1^ association with MTs (Ellefson & McNally, 2011; Crowder *et al*, 2015). The highest density of basic residues is found between residues 140-169 (30% K or R, pI = 12.02). This region is encoded by exon 4 and part of exon 5, which are subject to alternative splicing (Fig. 2B; S2A). This is strikingly similar to human p150, which contains an alternatively-spliced basic patch of 28 residues (43% K or R, pI = 12.7) adjacent to the CAP-Gly domain (Dixit *et al*, 2008). We detected four splice isoforms of *p150*^*dnc-1*^ by reverse transcription PCR of RNA isolated from adult animals (Fig. S2B): full-length (*FL*) *p150*^*dnc-1*^ including exons 4 and 5, *p150*^*dnc-1*^ without exon 4 (Δ*exon4*), *p150*^*dnc-1*^ without exon 5 (Δ*exon5*), and *p150*^*dnc-1*^ lacking exons 4 and 5 (Δ*exon4-5*). To define the function of individual splice isoforms, we edited the *p150*^*dnc-1*^ locus to generate animals in which *p150*^*dnc-1*^ expression was restricted to one of the four isoforms (Fig. 2B; S2A). Reverse transcription PCR confirmed that animals expressed single *p150*^*dnc-1*^ isoforms corresponding to *FL*, Δ*exon4*, Δ*exon5*, or Δ*exon4-5* (Fig. S2B). All mutant animals were homozygous viable and fertile (Fig. S2C), demonstrating that none of the p150^DNC-1^ splice isoforms is essential. Despite differences in predicted molecular weight of only a few kDa (Fig. S2A), single isoforms expressed in mutant animals were distinguishable by size on immunoblots with an antibody raised against a C-terminal region of p150^DNC-1^ (Fig. 2C). Side-by-side comparison of isoform mutants and wild-type animals on the same immunoblot revealed that neither the FL nor the Δexon4-5 isoform is prevalent in wild-type adults (Fig. 2C). Instead, immunoblotting, together with reverse transcription PCR data (Fig. S2B, D), suggested that p150^DNC-1^ Δexon4 is the predominant p150^DNC-1^ isoform.

**Figure 2:**
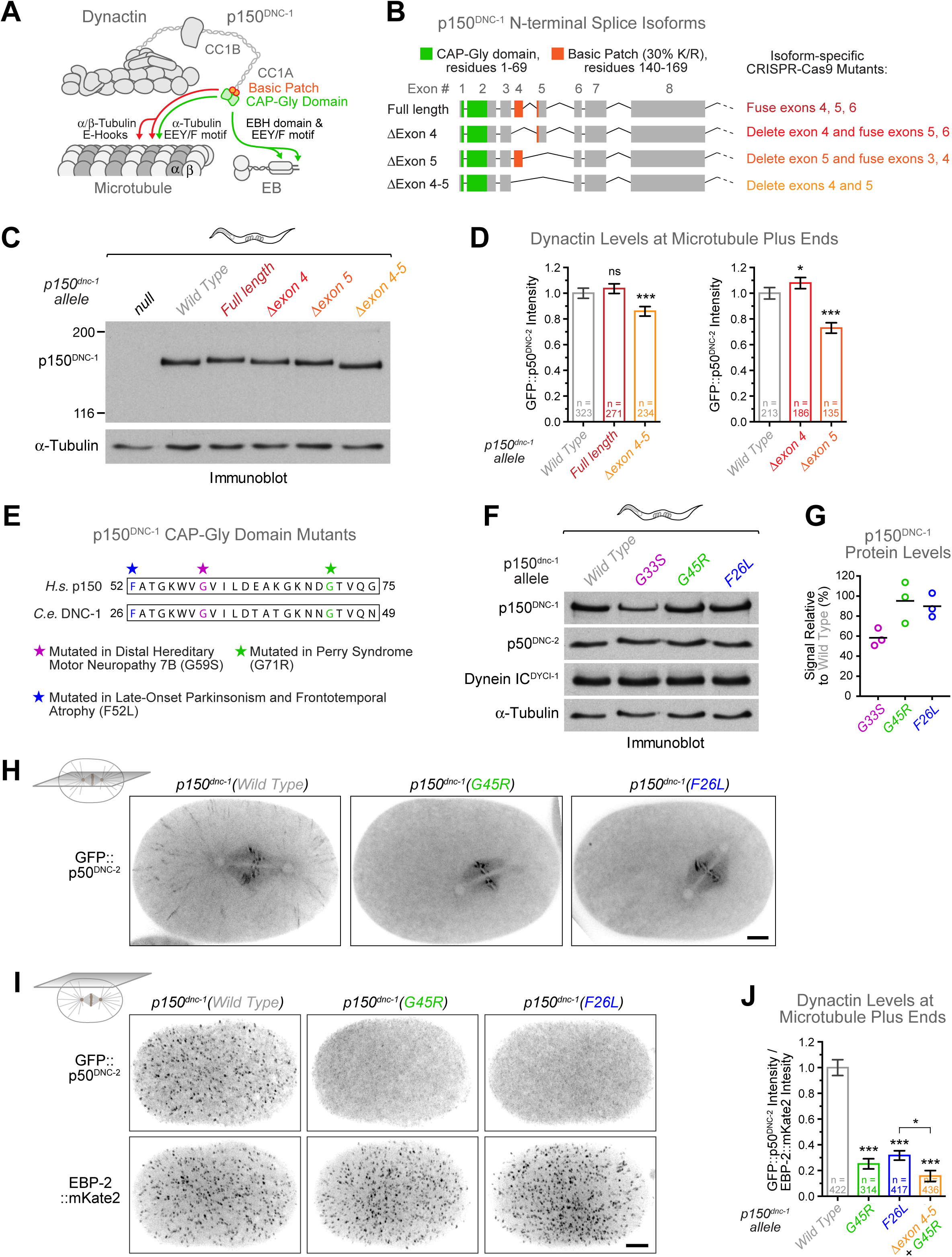
Engineering of p150^DNC-1^ mutants for functional characterization of dynactin’s MT binding region. **(A)** Cartoon of the dynactin complex and its interaction with MTs and +TIPs. The conserved N-terminal region of the p150^DNC-1^ subunit contains a CAP-Gly domain (residues 1-69) predicted to bind the end-binding homology (EBH) domain of end-binding proteins (EBs) and the EEY/F motifs present at the C-termini of EBs and α-tubulin. It also contains a basic patch (residues 140-169) predicted to interact with the negatively charged C-terminal tails (E-Hooks) of α- and β-tubulin. **(B)** Schematic of the four *p150*^*dnc-1*^ N-terminal splice isoforms identified by RT-PCR and strategy for CRISPR-Cas9-based genome editing to restrict *p150*^*dnc-1*^ expression to single isoforms. **(C)** Immunoblot of adults worms with an antibody against a C-terminal region of p150^DNC-^ 1, showing that engineered p150^dnc-1^ mutants express single isoforms that are distinguishable by size. α-Tubulin was used as loading control. Molecular mass is indicated in kilodaltons. **(D)** Quantification of GFP::p50^DNC-2^ levels at MT plus ends in controls and p150^DNC-1^ isoform mutants using fluorescence intensity measurements at the cortex. Error bars represent the SEM with a 95 % confidence interval, and *n* indicates the total number of individual MT plus ends measured in 5-8 different embryos per condition. The t-test was used to determine statistical significance (*** indicates p < 0.0001; * indicates p < 0.05; ns = not significant, p > 0.05). **(E)** Sequence alignment of p150 CAP-Gly domain region where point mutations have been identified that cause neurodegenerative disease in humans. Analogous point mutations were introduced into the *C. elegans p150*^*dnc-1*^ locus using CRISPR-Cas9-based genome editing. **(F)** Immunoblots of adult worms, showing that protein levels of p150^DNC-1^ are decreased in the G33S mutant but not in the G45R and F26L mutant. α-Tubulin served as the loading control. **(G)** Quantification of p150^DNC-1^ protein levels using intensity measurements from immunoblots as in *(F)*. Three independent immunoblots were performed for each condition. **(H)** Central confocal section in metaphase one-cell embryos expressing GFP::p50^DNC-2^ and wild-type p150^DNC-1^ or the G45R and F26L mutants. Image corresponds to a maximum intensity projection over time (10 images acquired every 200 ms). Scale bar, 5 *μ*m. **(I)** Cortical confocal section of embryos as in *(H)*, additionally expressing EBP-2::mKate2 as a marker for MT plus ends. Image corresponds to a maximum intensity projection over time (12 images acquired every 5 s). Scale bar, 5 *μ*m. **(J)** Quantification of dynactin levels at MT plus ends using fluorescence intensity measurements of GFP::p50^DNC-2^ at the cortex, as shown in *(I)*. For each MT end, GFP::p50^DNC-2^ signal was normalized to EBP-2::mKate2 signal. Error bars represent the SEM with a 95 % confidence interval, and *n* indicates the number of individual MT ends analyzed from 8 different embryos per condition. The t-test was used to determine statistical significance (*** indicates p < 0.0001; * indicates p < 0.05).

Humans express the neuron-specific splice variant p135, which lacks the entire N-terminal MT binding region (Tokito *et al*, 1996). In *C. elegans* hermaphrodite adults, 302 out of 959 somatic cells are neurons, yet we did not find evidence for a p135 isoform at the mRNA level (Fig. S2D), nor did our p150^DNC-1^ antibody detect proteins below ∼150 kDa in wild-type animals (Fig. 2C). We also generated a *p150*^*dnc-1*^*::3xflag* knock-in allele, and immunoblotting with antibody against 3xFLAG similarly failed to detect a p135 isoform (Fig. S2E). We speculated that specifically suppressing the expression of p150^DNC-1^ isoforms might facilitate the detection of p135 and engineered a null allele of *p150*^*dnc-1*^ by inserting a stop codon in exon 1 immediately following the start codon (Fig. S2A). The null mutation did not affect splicing of *p150*^*dnc-1*^ mRNA (Fig. S2B) and therefore should permit expression of a p135 isoform from an alternative start codon, as is the case in humans (Tokito *et al*, 1996). However, immunoblotting produced no evidence of p135 expression in the absence of p150^DNC-1^ isoforms (Fig. 2C). We conclude that *C. elegans*, in contrast to other species including vertebrates and *D. melanogaster*, does not express significant amounts of a p135 isoform.

We used cortical imaging of GFP::p50^DNC-2^ in one-cell embryos to determine the effect of p150^DNC-1^ isoform mutants on dynactin recruitment to MT tips. p150^DNC-1^ FL and Δexon4 fully supported dynactin targeting to MT tips, and dynactin levels were even slightly increased (108 ± 4 % of controls) for the Δexon4 isoform (Fig. 2D). By contrast, expression of p150^DNC-1^ Δexon5 or Δexon4-5 decreased dynactin levels at MT tips to 73 ± 4 % and 86 ± 4 % of controls, respectively. Thus, surprisingly, the similarly basic regions encoded by exon 4 (27% K/R; pI = 11.2) and exon 5 (19% K/R; pI = 12) make differential contributions to dynactin targeting. We conclude that p150^DNC-1^ splice isoforms regulate dynactin levels at MT plus ends.

### Point mutations in p150^DNC-1^’s CAP-Gly domain that cause neurodegenerative disease in humans displace dynactin and dynein from microtubule plus ends

Our analysis of p150^DNC-1^ isoform mutants suggested that dynactin targeting to MT tips primarily depended on the CAP-Gly domain rather than the adjacent basic region. To test this idea directly, we used genome editing to separately introduce three point mutations into p150^DNC-1^ that compromise CAP-Gly domain function and cause neurodegenerative disease in humans (Fig. 2E): G33S corresponds to human G59S, which causes motor neuropathy 7B (Puls *et al*, 2003); G45R corresponds to human G71R, which causes Perry Syndrome (Farrer *et al*, 2009); and F26L corresponds to human F52L, which was recently identified in a patient with Perry Syndrome-like symptoms (Araki *et al*, 2014). For the F26L and G45R mutations, animals could be propagated as homozygous mutants with high embryonic viability (99 ± 1 % and 90 ± 2 %, respectively), whereas the G33S mutant was lethal in the F2 generation (4.62 ± 3.75 % embryonic viability) (Fig. S3A). Immunoblotting of homozygous F1 adults showed that G33S animals had decreased levels of p150^DNC-1^, indicating that the mutation destabilized the protein (Fig. 2F, G). By contrast, total levels of p150^DNC-1^ were not affected by the F26L or G45R mutation. Central plane imaging in one-cell embryos expressing GFP::p50^DNC-2^ revealed that dynactin containing p150^DNC-1^ F26L or G45R was present on the mitotic spindle and prometaphase kinetochores but displaced from MT tips (Fig. 2H; Movies S3, S4). Cortical imaging after introduction of the EBP-2::mKate2 marker revealed that GFP::p50^DNC-2^ levels at MT tips were reduced to 32 ± 4 % and 25 ± 4 % of controls in the F26L and G45R mutant, respectively (Fig. 2I, J). Deletion of the basic patch encoded by exons 4 and 5 in the G45R mutant (Δexon4-5 + G45R) further reduced GFP::p50^DNC-2^ levels at MT tips to 16 ± 4 % (Fig. 2J). Additional quantifications showed that in both the F26L and G45R mutant, GFP::p50^DNC-2^ still targeted to the nuclear envelope and kinetochores, while GFP::p50^DNC-2^ levels were reduced on spindle MTs (Fig. S3B). We also introduced the p150^DNC-1^ F26L and G45R mutations into animals expressing DHC-1::GFP, which confirmed that dynein levels were decreased at MT tips and on spindle MTs (Fig. S4). We conclude that point mutations in the p150^DNC-1^ CAP-Gly domain that cause human neurodegenerative disease reduce dynein-dynactin levels on MTs and greatly diminish the ability of dynein-dynactin to track with MT tips.

### MT tip tracking of dynein-dynactin is dispensable for targeting the motor to the cell cortex

One potential role for MT tip tracking of dynein-dynactin in animal cells is delivery of the motor to the cell cortex for off-loading to anchor proteins. To examine whether displacement from MT tips affected dynein-dynactin accumulation at the cell cortex, we imaged the 4-cell embryo, in which dynactin and dynein become prominently enriched at the EMS-P2 cell border prior to EMS and P2 spindle rotation (Zhang *et al*, 2008; Walston & Hardin, 2006). Quantification of GFP::p50^DNC-2^ and DHC-1::GFP levels at the EMS-P2 cell border revealed that cortical levels of dynein-dynactin were unchanged in the p150^DNC-1^ G45R mutant (Fig. S3C). We conclude that MT tip tracking of dynein-dynactin is dispensable for cortical localization of the motor in the *C. elegans* early embryo.

### p150^DNC-1^ CAP-Gly domain mutants exhibit defects in centration/rotation of the nucleus-centrosome complex, chromosome congression, and spindle rocking

Next, we asked whether our p150^DNC-1^ mutants affected dynein-dynactin function in the one-cell embryo. We crossed the mutants with animals co-expressing GFP::histone H2B and GFP::γ-tubulin, which allowed precise tracking of pronuclei and centrosomes, respectively (Fig. 3A). None of the mutants exhibited defects in centrosome separation, and pronuclear migration along the anterior-posterior axis proceeded with normal kinetics until pronuclear meeting, which occurred at the correct position in the posterior half of the embryo (Fig; 3B; S5A). However, subsequent centration of the nucleus-centrosome complex (NCC) slowed substantially in p150^DNC-1^ F26L, G45R, and G45R+Δexon4-5 mutants, and NCC rotation was defective (Fig. 3A - D; Movie S5). NCC centration was not significantly perturbed in the isoform mutants (Fig. S5A, B), but the FL and Δexon4-5 mutant exhibited defects in NCC rotation (Fig. 3D; S5C). In all mutants, spindle orientation recovered during prometaphase, so that the spindle axis was largely aligned with the anterior-posterior axis of the embryo at the time of anaphase onset (Fig. 3D; S5C).

**Figure 3:**
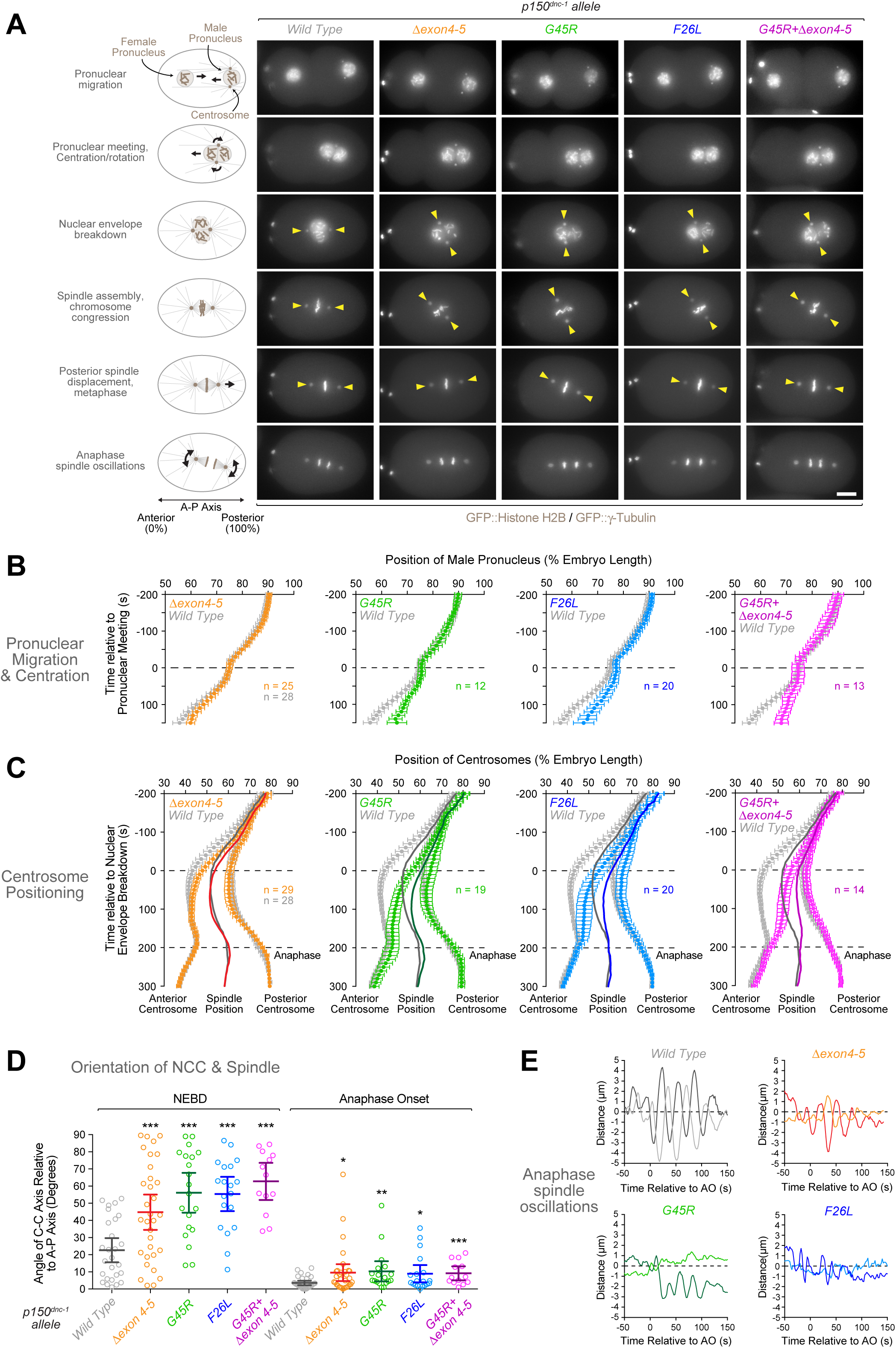
Mutations in the p150^DNC-1^ CAP-Gly domain and basic region cause defects in centration/rotation of the nucleus-centrosome complex. **(A)** Selected frames from time-lapse sequences of the first embryonic division in controls and p150^DNC-1^ mutants. Chromosomes and centrosomes are marked with GFP::histone H2B and GFP::γ-tubulin, respectively. Scale bar, 5 *μ*m. **(B)** Migration kinetics of the male pronucleus in the embryos shown in *(A)*. The position of the male pronucleus along the anterior-posterior axis was determined in images captured every 10 s. Individual traces were normalized to embryo length, time-aligned relative to pronuclear meeting, averaged for the indicated number (n) of embryos, and plotted against time. Error bars represent the SEM with a 95 % confidence interval. **(C)** Kinetics of centrosome positioning along the anterior-posterior axis, determined as described for *(B)* and plotted relative to NEBD. Solid lines indicate the midpoint between the two centrosomes (spindle position). Anaphase begins at 200 s. Error bars represent the SEM with a 95 % confidence interval. **(D)** Angle between the centrosome-centrosome (C-C) axis and the anterior-posterior (A-P) axis in one-cell embryos at NEBD and anaphase onset. Circles correspond to measurements in individual embryos. Error bars represent the SEM with a 95 % confidence interval. The t-test was used to determine statistical significance (*** indicates p < 0.0001; ** indicates p < 0.01; * indicates p < 0.05; ns = not significant, p > 0.05). **(E)** Representative examples of spindle transversal oscillation in anaphase. The **(A)** transverse position of spindle poles was determined every 2 s and plotted against time.

In controls, the mitotic spindle was displaced from the embryo center toward the posterior in preparation for asymmetric division (Fig. 3C). By contrast, spindle assembly in CAP-Gly domain mutants already occurred in the posterior half of the embryo, and the spindle had to be moved only slightly to the posterior to be correctly positioned. In controls, the regular and vigorous oscillations of spindle rocking began at anaphase onset and lasted for approximately 100 s. By contrast, spindle rocking in p150^DNC-1^ CAP-Gly mutants was irregular and significantly dampened (Fig. 3E; Movie S6).

In addition to defects in NCC centration/rotation and spindle rocking, we observed a slight but consistent delay in chromosome congression in p150^DNC-1^ CAP-Gly mutants, indicating problems with the interaction between chromosomes and spindle MTs (Fig. S6A, B; Movie S6). This did not result in obvious chromosome mis-segregation in the first embryonic division. However, when the spindle assembly checkpoint (SAC) was inactivated by RNA-mediated depletion of Mad1^MDF-1^, embryonic viability decreased by 28% and 22% in the G45R and F26L mutant, respectively, whereas *Mad1*^*MDF-1*^*(RNAi)* in controls decreased embryonic viability by just 6% (Fig. S6C). This suggests that SAC signaling is required during embryogenesis to prevent chromosome segregation errors when p150^DNC-1^ CAP-Gly domain function is compromised.

We conclude that mutations in the p150^DNC-1^ CAP-Gly domain perturb a specific subset of dynein-dynactin functions in the one-cell embryo.

### Inhibition of cortical dynein and p150^DNC-1^ CAP-Gly mutants cause distinct defects

Anaphase spindle rocking requires cortical dynein pulling on astral MTs. Since spindle rocking was affected in p150^DNC-1^ CAP-Gly mutants, we wanted to assess the extent of phenotypic overlap between p150^DNC-1^ CAP-Gly mutants and inhibition of dynein-dependent cortical pulling. We therefore tracked centrosomes and pronuclei after co-depleting GPR-1 and GPR-2, which are required for cortical anchoring of dynein-dynactin (Couwenbergs *et al*, 2007; Nguyen-Ngoc *et al*, 2007). In contrast to p150^DNC-1^ mutants, *gpr-1/2(RNAi)* delayed the initial separation of centrosomes and the onset of pronuclear migration (Fig. S7A, D). Pronuclear migration and NCC centration subsequently occurred at a slightly faster rate than in controls, so that the NCC achieved near-normal centration by NEBD (Fig. S7A, B). These results are consistent with slowed centrosome separation and faster centering reported after co-depletion of GOA-1 and GPA-16, the Gα proteins acting upstream of GPR-1/2 (Kimura & Onami, 2007; De Simone *et al*, 2016). Thus, the kinetics of pronuclear migration and NCC centration differ between *gpr-1/2(RNAi)* and p150^DNC-1^ CAP-Gly mutants. NCC rotation, by contrast, was affected in both perturbations. Importantly, *gpr-1/2(RNAi)* in the p150^DNC-1^ G45R mutant enhanced the rotation defect, arguing that GPR-1/2 and the p150^DNC-1^ CAP-Gly domain contribute to NCC rotation through parallel pathways (Fig. S7C). After NEBD, depletion of GPR-1/2 prevented posterior displacement of the spindle and the lack of cortical pulling was especially evident in the track of the posterior centrosome (Fig. S7B). In addition, the mitotic spindle was shorter than controls during metaphase and failed to elongate properly in anaphase (Fig. S7D). By contrast, posterior centrosome movement towards the cortex in p150^DNC-1^ CAP-Gly mutants was indistinguishable from controls (Fig. 3C; S7B), and spindle length was normal throughout metaphase and anaphase (Fig. S7D). These results argue that, although spindle rocking is compromised in p150^DNC-1^ CAP-Gly mutants, cortical dynein is still able to generate significant pulling forces. Of note, prior work suggested that anaphase spindle rocking is particularly sensitive to changes in parameters that affect cortical force generation, including MT dynamics (Pecreaux *et al*, 2006; Schmidt *et al*, 2005; Kozlowski *et al*, 2007). We did observe slight changes in the cortical residency time of EBP-2::mKate2 in p150^DNC-1^ CAP-Gly mutants (data not shown), which may provide an explanation for the spindle rocking defects.

In conclusion, our comparative analysis with *gpr-1/2(RNAi)* argues that the defects in NCC centration/rotation are unlikely to be caused by a lack of cortical pulling forces.

### MT tip tracking of dynein-dynactin is dispensable for cortical force generation

We showed that delocalization of dynein-dynactin from MT tips in p150^DNC-1^ CAP-Gly mutants did not prevent cortical targeting of the motor (Fig. S3C), nor did it prevent cortical dynein from generating robust pulling forces (Fig. S7A-D). To obtain further evidence that MT tip tracking of dynein-dynactin is dispensable for cortical force generation, we tracked centrosomes after depletion of EBP-2, which, just like p150^DNC-1^ CAP-Gly mutants, delocalizes dynein-dynactin from MT tips (Fig. 1B). Strikingly, posterior spindle displacement was exaggerated in *ebp-2(RNAi*) embryos compared with controls (Fig. S7B), and anaphase spindle rocking was normal (data not shown). This demonstrates that cortical pulling forces can be generated in the absence of MT tip-localized dynein-dynactin.

### Mutation of α-tubulin’s C-terminal tyrosine causes defects in NCC centration and rotation that resemble those of p150^DNC-1^ CAP-Gly domain mutants

CAP-Gly domains bind the C-terminal EEY/F motif of α-tubulin, and the tyrosine residue is critical for the interaction (Fig. 2A) (Steinmetz & Akhmanova, 2008). We therefore asked whether decreased affinity of dynactin for tyrosinated MTs could be contributing to the defects observed in p150^DNC-1^ CAP-Gly mutants. Of the 9 α-tubulin isoforms in *C. elegans*, *tba-1* and *tba-2* are the major α-tubulin isotypes expressed during early embryogenesis (Baugh *et al*, 2003). We mutated the C-terminal tyrosine of TBA-1 and TBA-2 to alanine and obtained animals homozygous for either mutation alone (YA) or both mutations combined (YA/YA) (Fig. 4A). Immunoblotting of adult animals with the monoclonal antibody Y1/2, which is specific for tyrosinated tubulin, revealed that levels of tubulin tyrosination were decreased in *tba-1(YA)* and *tba-2(YA)* single mutants, with *tba-2(YA)* having a more pronounced effect (Fig. 4B). Combining the two mutations dramatically decreased total levels of tubulin tyrosination. Importantly, immunoblotting with an antibody insensitive to tubulin tyrosination confirmed that total α-tubulin levels were not affected in the three mutants (Fig. 4B). We then used immunofluorescence to directly assess tyrosinated tubulin levels in the early embryo. In controls, the mitotic spindle of the one-cell embryo was prominently stained with the antibody against tyrosinated tubulin (Fig. 4C). By contrast, the tubulin tyrosination signal was undetectable in the *tba-1/2(YA/YA)* double mutant, despite normal spindle assembly. Thus, we generated animals without detectable tubulin tyrosination in early embryos.

**Figure 4:**
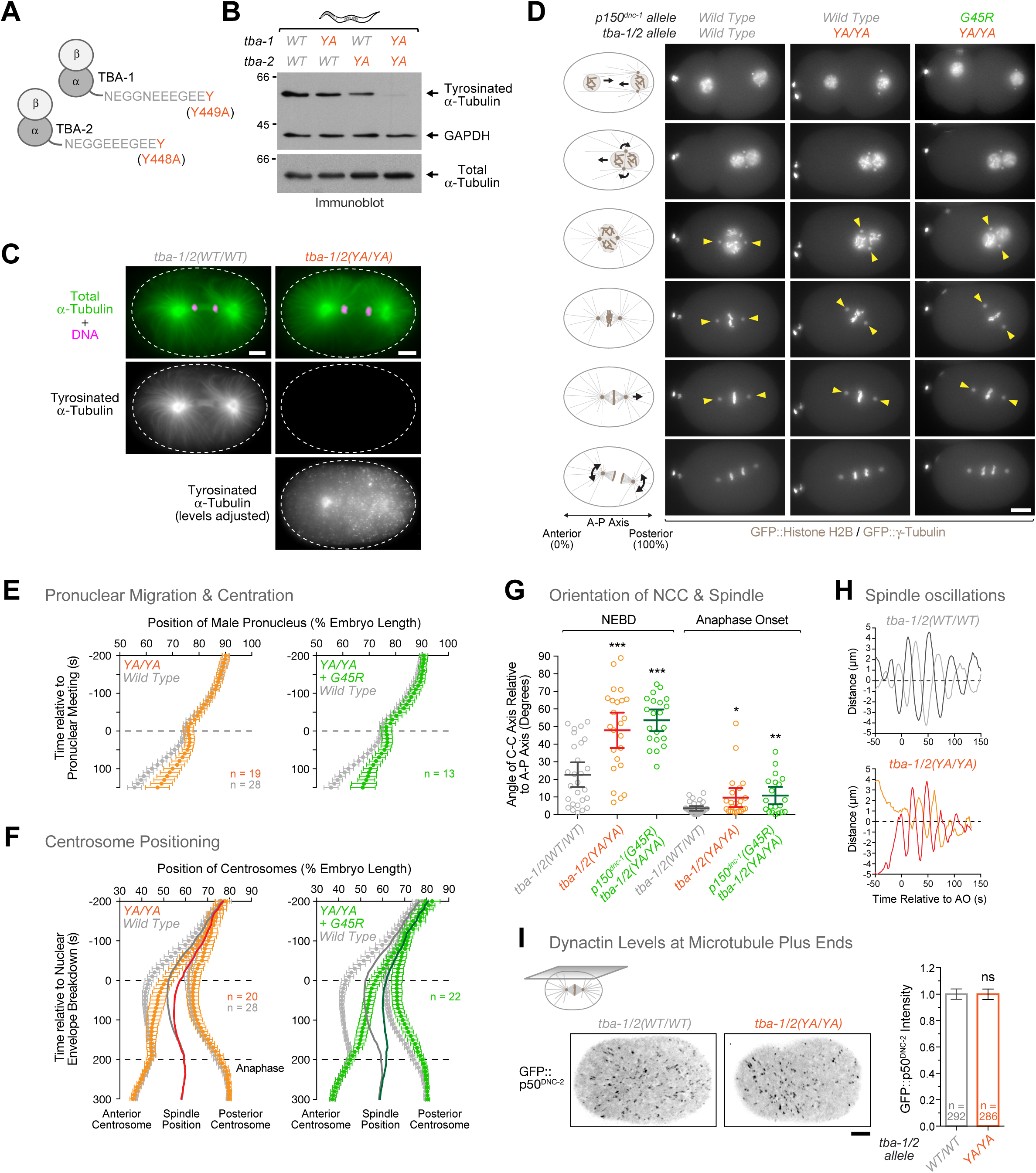
The C-terminal tyrosine of α-tubulin is required for centration/rotation of the nucleus-centrosome complex but not for MT plus-end targeting of dynactin. **(A)** Cartoon depicting the C-terminal tail of TBA-1 and TBA-2, the major α-tubulin isotypes in early embryogenesis. The C-terminal tyrosine residues were mutated to alanine (YA) using CRISPR-Cas9-based genome editing. **(B)** Immunoblot of adult worms showing decreased levels of tubulin tyrosination in α-tubulin YA mutants. The same membrane was probed sequentially with an antibody against tyrosinated α-tubulin (monoclonal antibody Y1/2) and against total α-tubulin (monoclonal antibody B512). GAPDH served as a loading control. Molecular mass is indicated in kilodaltons. **(C)** Immunofluorescence images showing lack of tubulin tyrosination in *tba-1/2(YA/YA)* embryos at the one-cell stage (anaphase). Embryos were co-stained with antibodies against tyrosinated α-tubulin and total α-tubulin. Image with adjusted intensity levels shows unspecific background signal in the α-tubulin mutant. Scale bar, 5 *μ*m. **(D) - (H)** Still images from time-lapse sequences *(D)* and analysis of male pronuclear migration *(E)*, centrosome position *(F)*, NCC rotation/spindle orientation *(G)*, and anaphase spindle oscillations *(H)* during the first embryonic division for controls and α-tubulin tyrosine mutants. Data is displayed as described for Figure 3. Scale bar, 5 *μ*m. **(I)** *(left)* Cortical confocal section of a control and *tba-1/2(YA/YA)* one-cell embryo in metaphase expressing GFP::p50^DNC-2^. Image corresponds to a maximum intensity projection over time of 12 images acquired every 5 s. Scale bar, 5 *μ*m. *(right)* Quantification of dynactin levels at MT plus ends using fluorescence intensity measurements of GFP::p50^DNC-2^ at the cortex. Error bars represent the SEM with a 95 % confidence interval, and *n* indicates the number of individual MT ends analyzed from 8-9 different embryos per condition. The t-test was used to determine statistical significance (ns = not significant, p > 0.05).

We then tested the effect of the *tba-1/2(YA/YA)* mutant in the one-cell embryo. Strikingly, we found that the *tba-1/2(YA/YA)* mutant exhibited NCC centration/rotation defects reminiscent of those observed in p150^DNC-1^ G45R and F26L mutants (Fig. 4D-G; Movie S7). Importantly, combining the *tba-1/2(YA/YA)* mutant with the p150^DNC-1^ G45R mutant did not exacerbate the centration/rotation defects of the p150^DNC-1^ G45R mutant alone, indicating that both mutants act in the same pathway. Like in p150^DNC-1^ CAP-Gly mutants, we noticed a slight delay in chromosome congression in the *tba-1/2(YA/YA)* mutant (data not shown), while anaphase spindle rocking was normal (Fig. 4H).

We also examined the effect of the *tba-1/2(YA/YA)* mutant on GFP::p50^DNC-2^ localization and found that dynactin levels at MT tips were identical to controls (Fig. 4I). Thus, in contrast to mouse fibroblasts (Peris *et al*, 2006), tubulin tyrosination in the *C. elegans* embryo is not required to target dynactin to MT tips.

### p150^DNC-1^ CAP-Gly mutants and α-tubulin tyrosine mutants decrease the frequency of centrosome-directed early endosome movements

Dynein-mediated transport of small organelles along MTs towards centrosomes is proposed to generate the cytoplasmic pulling forces for centration (the centrosome-organelle mutual pulling model) (Kimura & Onami, 2005; Kimura & Kimura, 2011; Shinar *et al*, 2011). To ask whether the centration defects in our mutants correlate with defects in MT minus-end-directed organelle transport, we monitored the movements of early endosomes, labelled with mCherry::RAB-5, from pronuclear meeting until nuclear envelope breakdown. Time-lapse sequences, recorded at 2.5 frames per second in a focal plane that included the NCC, were used for semi-automated tracking of early endosomes that moved from the cell periphery towards centrosomes (Fig. 5A). In control embryos, we counted 16.3 ± 3.5 tracks/min during the ∼6 min interval of centration (Fig. 5B). This was reduced to 0.8 ± 0.6 tracks/min in embryos depleted of p150^DNC-1^ by RNAi, confirming that dynactin is required for early endosome movement directed towards centrosomes. The p150^DNC-1^ G45R+Δexon4-5 mutant also strongly reduced the number of observed tracks to 5.3 ± 1.1 tracks/min (Fig. 5B; Movie S8). The TBA-1/2(YA/YA) mutant had a less severe effect but still substantially reduced the number of tracks to 10.6 ± 2.2 per min. We also determined the maximal velocity in each track (determining the mean speed was complicated by frequent pausing of particles) and the total track displacement. This revealed only minor differences between controls and either mutant (Fig. 5B). We conclude that p150^DNC-1^ CAP-Gly and α-tubulin tyrosine mutants reduce the frequency with which early endosomes move towards centrosomes but not their maximal velocity or the overall length of the distance travelled. These results are consistent with the idea that dynactin binding to tyrosinated MTs enhances the efficiency of transport initiation by dynein, as recently documented *in vitro* (McKenney *et al*, 2016).

**Figure 5:**
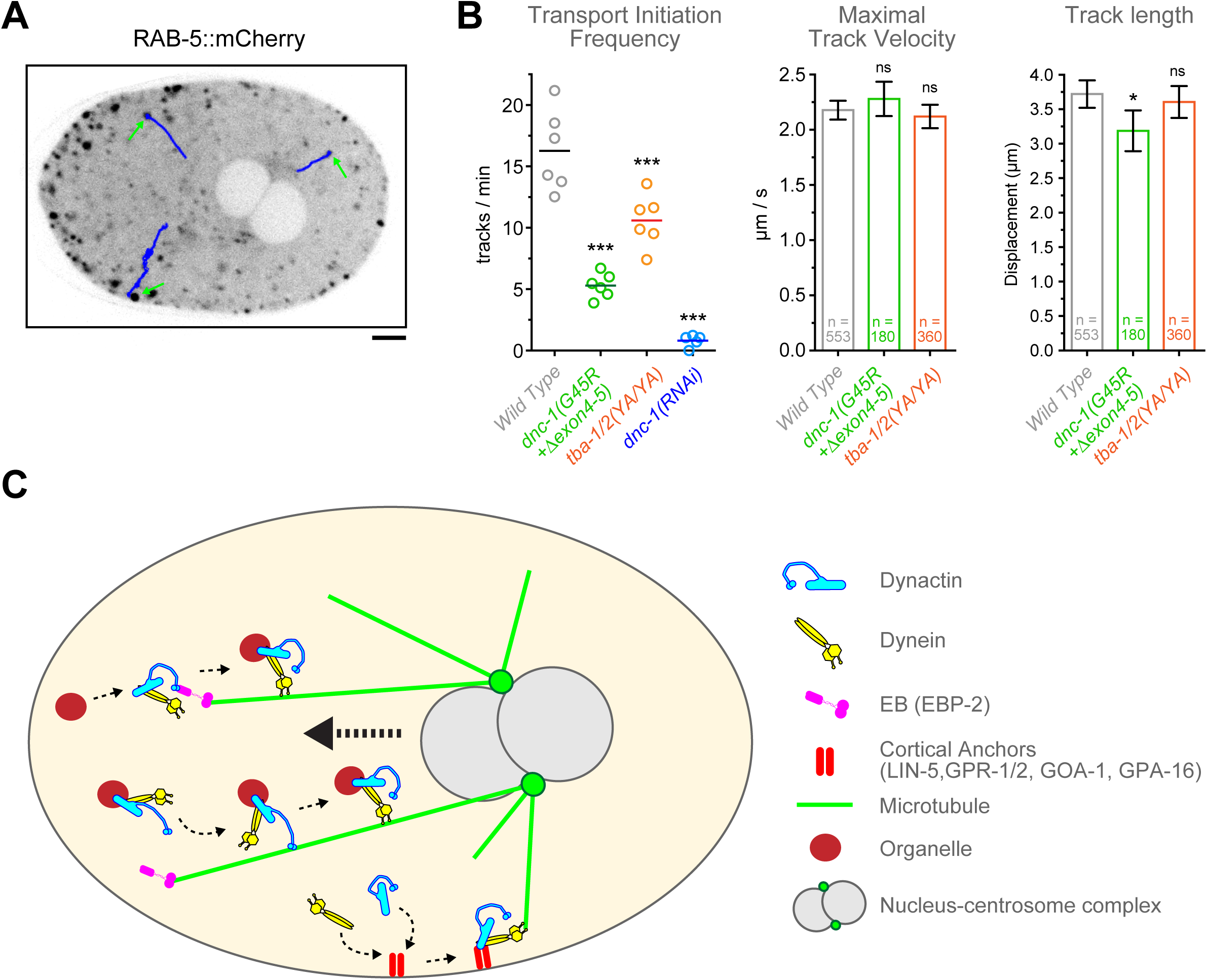
Dynactin binding to MTs and tyrosinated α-tubulin are required for efficient initiation of centrosome-directed organelle transport. **(A)** Selected image from a time-lapse sequence recorded during centration/rotation of the nucleus-centrosome complex in an embryo expressing the early endosome marker mCherry::RAB-5. Arrows point to early endosomes that are about to move from the cell periphery towards the centrosomes along the tracks shown in blue. **(B)** Quantification of the number of tracks per min, the maximal track velocities, and the total track displacement for mCherry::RAB-5-labeled particles travelling towards centrosomes during the centration/rotation phase. Particles were tracked in time-lapse sequences as shown in *(A)* with images captured every 400 ms. Circles correspond to data points of individual embryos. Error bars represent the SEM with a 95 % confidence interval, and *n* indicates the number of individual tracks analyzed from 8-9 embryos per condition. The t-test was used to determine statistical significance (*** indicates p < 0.0001; ** indicates p < 0.01; * indicates p < 0.05; ns = not significant, p > 0.05). **(C)** Cartoon model summarizing the functional significance of dynactin’s MT binding activity in the one-cell embryo. The CAP-Gly domain of p150^DNC-1^ binds to MTs and MT tips through an interaction with the C-terminal tyrosine of α-tubulin and EBP-2, respectively. This promotes the initiation of organelle transport by dynein from MT tips and along the length of MTs. Dynein-dependent movement of organelles along MTs towards centrosomes generates pulling forces required for centration of the nucleus-centrosome complex. In p150^DNC-1^ CAP-Gly mutants and α-tubulin tyrosine mutants, the initiation of organelle transport occurs with lower efficiency, which decreases cytoplasmic pulling forces and therefore delays centration. Recruitment of dynein-dynactin to MT tips is not required for targeting dynein to cortical anchors. Instead, dynein-dynactin bind to the cortex directly from the cytoplasm and subsequently capture MT tips for generation of cortical pulling forces.

## DISCUSSION

Dynactin’s MT binding activity is crucial in neurons, as illustrated by single point mutations that compromise the function of the p150 CAP-Gly domain and cause neurodegenerative disease (Lenz *et al*, 2006; Lloyd *et al*, 2012; Moughamian & Holzbaur, 2012; Zhang *et al*, 2010). Here, we introduced these CAP-Gly mutations into *C. elegans* p150^DNC-1^ to investigate how dynactin’s interaction with MTs and +TIPs contributes to dynein function in early embryogenesis. Together with the analysis of engineered p150^DNC-1^ splice isoforms and tubulin tyrosination mutants, our work provides insight into the regulation and function of MT tip tracking by dynein-dynactin in metazoans and uncovers a link between dynactin’s role in initiating dynein-mediated transport of small organelles and the generation of cytoplasmic pulling forces.

Dynein accumulates at and tracks with growing MT plus ends in species ranging from fungi to mammals, but requirements for MT tip tracking differ. In the *C. elegans* early embryo, MT tip recruitment of dynein-dynactin shares similarity with the pathway in budding yeast (dynactin depends on dynein and LIS-1) and mammalian cells/filamentous fungi (dynein depends on dynactin). Surprisingly, similar to what was reported for the fungus *U. maydis* (Lenz *et al*, 2006), accumulation of dynein-dynactin at MT tips does not require a CLIP-170-like protein in *C. elegans*. Instead, dynactin is likely directly recruited by EBP-2, one of the three EB homologs. Work in mouse fibroblasts knocked out for tubulin tyrosine ligase showed that decreased tyrosinated tubulin levels displaced CLIP-170 and p150 from MT plus ends (Peris *et al*, 2006). By contrast, we show that MT tip targeting of *C. elegans* dynactin is independent of tubulin tyrosination, possibly because there is no requirement for a CLIP-170 homolog. Overall, our analysis of dynein-dynactin targeting to MT tips in *C. elegans* highlights the evolutionary plasticity of +TIP networks.

Our analysis of engineered p150^DNC-1^ mutants establishes the functional hierarchy among p150’s tandem arrangement of MT binding regions: the CAP-Gly domain clearly provides the main activity, while the adjacent basic region plays an auxiliary role. Together with previous work in human cells (Dixit *et al*, 2008), our results support the idea that alternative splicing of p150’s basic region constitutes a conserved mechanism in metazoans for fine-tuning dynactin’s affinity for MTs. Only one of the four p150^DNC-1^ splice isoforms we identified, p150^DNC-1^ Δexon4, fully supported dynactin localization and function in the one-cell embryo, suggesting that the basic region has cell type-specific roles.

In budding yeast, dynein must first be targeted to MT tips prior to associating with cortical anchors (Markus & Lee, 2011b; 2011a). Our results suggest that this pathway is not used in *C. elegans*, as cortical accumulation of dynein-dynactin was unaffected in the p150^DNC-1^ G45R mutant that displaced the majority of dynactin and dynein from MT tips. In agreement with normal cortical targeting of the motor, dynein-dependent cortical pulling forces were not significantly affected in p150^DNC-1^ CAP-Gly mutants, although the defects in spindle rocking shows that the p150^DNC-1^ CAP-Gly domain does contribute to cortical force generation in anaphase. Importantly, delocalizing dynein-dynactin from MT tips by depletion of EBP-2 even enhanced cortical pulling during posterior spindle displacement. Thus, our results argue that dynein is recruited by cortical anchors directly from the cytoplasm, and that dynein-dependent cortical force generation is therefore functionally uncoupled from MT tip tracking of the motor (Fig. 5C). It will be interesting to determine whether that is also the case in other animals.

If not delivery of the motor to the cell cortex, what is the purpose of dynein’s MT tip tracking behavior in the embryo? We found that p150 CAP-Gly mutants have defects in the centration and rotation of the NCC, which consists of the two centrosomes and the associated female and male pronucleus. Experimental work and biophysical modelling support the idea that centration forces in the one-cell embryo are generated by dynein-mediated cytoplasmic pulling (Kimura & Onami, 2005; Kimura & Kimura, 2011; Shinar *et al*, 2011), although a centration/rotation model based on cortical pulling forces has also been proposed (Coffman *et al*, 2016). In the cytoplasmic pulling model, dynein works against viscous drag as it transports small organelles (e.g. endosomes, lysosomes, yolk granules) along MTs towards centrosomes, which generates pulling forces on MTs that move the NCC. Prior work showed that movements of early endosomes and centrosomes are correlated, and RNAi-mediated depletion of proteins that tether dynein to early endosomes and lysosomes inhibited centration, indicating that there is a functional link between organelle transport and cytoplasmic pulling forces (Kimura & Kimura, 2011). In agreement with this idea, the p150^DNC-1^ G45R+Δexon4-5 mutant not only inhibited centration but also significantly decreased the number of early endosomes that displayed directed movement toward centrosomes. This effect on early endosome transport is consistent with the p150 CAP-Gly domain’s role in initiating dynein-mediated transport, which is well-established in the context of retrograde axonal transport in neurons (Lloyd *et al*, 2012; Moughamian & Holzbaur, 2012; Moughamian *et al*, 2013). Compromising the efficiency with which organelle transport is initiated is predicted to decrease cytoplasmic pulling forces, because the magnitude of the net pulling force acting on centrosomes is proportional to the number of organelles travelling along MTs.

The frequency of centrosome-directed early endosome movement was also decreased in the TBA-1/2(YA/YA) mutant, which severely reduces the levels of tubulin tyrosination in the early embryo. This fits well with recent work *in vitro* demonstrating that the interaction between the p150 CAP-Gly domain and tyrosinated MTs enhances the efficiency with which processive motility of dynein-dynactin is initiated (McKenney *et al*, 2016). Furthermore, a recent study in neurons provided evidence that initiation of retrograde transport in the distal axon is regulated by α-tubulin tyrosination (Nirschl *et al*, 2016).

Strikingly, the tyrosination mutant TBA-1/2(YA/YA) also affected centration of the NCC, as predicted by the centrosome-organelle mutual pulling model. Of note, the effect on early endosome transport was slightly more pronounced in the p150^DNC-1^ G45R+Δexon4-5 mutant than in the tubulin tyrosine mutant, and this correlated with a more pronounced defect in centration/rotation. The difference could be explained by imagining that transport initiation can occur both from the MT tip (facilitated by the pool of dynein-dynactin localized there through the interaction between the p150^DNC-1^ CAP-Gly domain and EBP-2) and along the entire length of the MT (facilitated by the interaction between the p150^DNC-1^ CAP-Gly domain and the C-terminal tyrosine of α-tubulin (McKenney *et al*, 2016). The p150^DNC-1^ G45R+Δexon4-5 mutant will inhibit both modes of transport initiation, whereas the tubulin tyrosine mutant does not affect dynein-dynactin recruitment by EBP-2 and may therefore still support transport initiation from MT tips (Fig.5C).

Why do the p150^DNC-1^ and α-tubulin mutants affect centration/rotation of the NCC, but not pronuclear migration until pronuclear meeting? One obvious explanation is that during pronuclear migration the male and female pronuclei, which are large (∼10 *μ*m diameter) and equal in size, assist each other’s movement as dyneins anchored on the female pronucleus walk along MTs nucleated by the centrosomes attached to the male pronucleus (Reinsch & Gönczy, 1998). By contrast, during centration, the two pronuclei must be moved in the same direction, which might render cytoplasmic pulling forces more sensitive to changes in the frequency of centrosome-directed organelle transport.

Finally, our data suggest that MT binding by dynactin contributes to chromosome congression. The effect is unlikely an indirect consequence of the delay in spindle orientation along the A-P axis, as alignment problems were not observed after *gpr-1/2(RNAi)*, which also causes spindle orientation defects. Likewise, normal chromosome congression in *ebp-2(RNAi)* suggests that the delay in chromosome congression in p150^DNC-1^ CAP-Gly mutants is not due to delocalization of dynactin from MT tips. Therefore, it is likely that the contribution to chromosome congression comes from the p150^DNC-1^ CAP-Gly domain pool at kinetochores, where it could aid in the capture of MTs. In support of this idea, the α-tubulin tyrosine mutant, which is predicted to affect MT binding to the CAP-Gly domain, also exhibited delayed chromosome congression (data not shown). The decrease in embryonic viability in p150^DNC-1^ CAP-Gly domain mutants after inhibition of Mad1^MDF-1^ indicates that chromosome congression problems persist in later embryonic divisions and compromise the fidelity of chromosome segregation.

In summary, our work demonstrates that dynactin’s MT binding activity is functionally relevant in the context of embryonic cell division. Unlike previous work that addressed CAP-Gly domain function in dividing S2 cells (Kim *et al*, 2007), we do not observe defects in bipolar spindle formation in p150^DNC-1^ CAP-Gly mutants. Instead, the most striking consequence of inhibiting p150^DNC-1^ CAP-Gly function or tubulin tyrosination is defective centrosome centration, which we propose is a consequence of defective initiation of dynein-mediated organelle transport, in agreement with the centrosome-organelle mutual pulling model. The transport initiation function of p150’s CAP-Gly domain is likely generally relevant in circumstances where positioning of subcellular structures depends on dynein-mediated cytoplasmic pulling, for example the centration of sperm asters in the large eggs of amphibians and sea urchins (Wühr *et al*, 2010; Tanimoto *et al*, 2016).

## MATERIALS AND METHODS

### Worm strains

Worm strains used in this study are listed in Table S1. Worms were maintained at 16, 20 or 25 °C on standard NGM plates seeded with OP50 bacteria. A Mos1 transposon-based strategy (MosSCI) was used to generate strains stably expressing EBP-2::mKate2 (Frøkjaer-Jensen *et al*, 2012). The transgene was cloned into pCFJ151 for insertion on chromosome II (ttTi5605 locus), and transgene integration was confirmed by PCR. The following alleles were generated by CRISPR-Cas9-based genome editing, as described previously (Arribere *et al*, 2014; Paix *et al*, 2014): *gfp::p50*^*dnc-2*^, *dynein heavy chain*^*dhc-*^*1::gfp*, *p150*^*dnc-1*^ *F26L*, *p150*^*dnc-1*^ *G33S*, *p150*^*dnc-1*^ *G45R*, *p150*^*dnc-1*^ *exon 4-5-6 fusion*, *p150*^*dnc-1*^ Δ*exon 4/exon 5-6 fusion*, *p150*^*dnc-1*^ Δ*exon 5/exon 3-4 fusion*, *p150*^*dnc-1*^ Δ*exon 4-5*, *p150*^*dnc-1*^ *null*, α*-tubulin*^*tba-1*^ *Y449A*, α*-tubulin*^*tba-2*^ *Y448A*, *CLIP-170*^*clip-1*^ *null*, and *p150*^*dnc-*^ *1::3xflag*. Genomic sequences targeted by the sgRNAs are listed in Table S2. The modifications were confirmed by sequencing and strains were outcrossed 6 times with the wild-type N2 strain. Other fluorescent markers were subsequently introduced by mating. The *p150*^*dnc-1*^ *G33S* allele and the *p150*^*dnc-1*^ *null* allele were maintained using the GFP-marked genetic balancer nT1 [qIs51]. Homozygous F1 progeny from balanced heterozygous mothers was identified by the lack of GFP fluorescence. None of the homozygous F1 *p150*^*dnc-1*^ *null* progeny reached adulthood, and homozygous F2 *p150*^*dnc-1*^ *G33S* progeny died during embryogenesis.

### RNA interference

For production of double-stranded RNAs (dsRNA), oligos with tails containing T3 and T7 promoters were used to amplify regions from N2 genomic DNA or cDNA. Primers are listed in Table S3. PCR reactions were cleaned (NucleoSpin® Gel and PCR Clean-up, Macherey-Nagel) and used as templates for T3 and T7 transcription reactions (MEGAscript®, Invitrogen). Transcription reactions were cleaned (NucleoSpin® RNA Clean-up, Macherey-Nagel) and complementary single-stranded RNAs were annealed in soaking buffer (3x soaking buffer is 32.7 mM Na_2_HPO_4_, 16.5 mM KH_2_PO_4_, 6.3 mM NaCl, 14.1 mM NH_4_Cl). dsRNAs were delivered by injecting L4 hermaphrodites, and animals were processed for live imaging after incubation at 20 °C for 24 or 48 h for partial and penetrant depletions, respectively.

### Antibodies

An affinity-purified rabbit polyclonal antibody against the N-terminal region of dynein intermediate chain^DYCI-1^ (residues 1-177) was generated as described previously (Desai *et al*, 2003). In brief, a GST fusion was expressed in *E. coli*, purified, and injected into rabbits. Serum was affinity purified on a HiTrap *N*-hydroxysuccinimide column (GE Healthcare) against covalently coupled DYCI-1_1-177_. The antibodies against p150^DNC-1^ (GC2) and p50^DNC-2^ (GC5) were described previously (Gama *et al*, 2017).

### Indirect immunofluorescence

For immunofluorescence of *C. elegans* embryos, 10 - 12 adult worms were dissected into 3 *μ*L of M9 buffer (86 mM NaCl, 42 mM Na_2_HPO_4_, 22 mM KH_2_PO_4_, 1 mM MgSO_4_) on poly-*L*-lysine-coated slides. A 13 mm^2^ round coverslip was placed on the 3 *μ*l drop, and slides were plunged into liquid nitrogen. After rapid removal of the coverslip (“freeze-cracking”), embryos were fixed in -20 °C methanol for 20 min. Embryos were re-hydrated for 2 × 5 min in PBS (137 mM NaCl, 2.7 mM KCl, 8.1 mM Na_2_HPO_4_, and 1.47 mM KH_2_PO_4_), blocked with AbDil (PBS with 2 % BSA, 0.1 % Triton X-100) in a humid chamber at room temperature for 30 minutes, and incubated with primary antibodies [mouse monoclonal anti-α-tubulin DM1 (1:1000) and rat monoclonal anti-tyrosinated α-tubulin YL1/2 (1:500)] for 2 h at room temperature. After washing for 4 × 5 min in PBS, embryos were incubated with secondary antibodies conjugated with fluorescent dyes (Alexa Fluor®488 goat anti-rat IgG (1:1000) and Alexa Fluor® 568 goat anti-mouse IgG (1:1000); Life Technologies - Molecular Probes) for 1h at room temperature. Embryos were washed for 4 × 5 min in PBS and mounted in Prolong® Gold with DAPI stain (Invitrogen).

Images were recorded on an inverted Zeiss® Axio Observer microscope, at 1 × 1 binning with a 100x NA 1.46 Plan-Apochromat objective and an Orca Flash 4.0 camera (Hamamatsu). Image files were imported into Fiji for further processing.

### Immunoblotting

For each condition, 100 worms were collected into 1 mL M9 buffer and washed 3 × with M9 buffer and once with M9 / 0.05 % Triton X-100. To a 100 *μ*L worm suspension, 33 *μ*L 4x SDS-PAGE sample buffer [250 mM Tris-HCl, pH 6.8, 30 % (v/v) glycerol, 8 % (w/v) SDS, 200 mM DTT and 0.04 % (w/v) bromophenol blue] and ∼20 *μ*L of glass beads were added. Samples were incubated for 3 min at 95 °C and vortexed for 2 × 5 min. After centrifugation at 20000 × g for 1min at room temperature, supernatants were collected and stored at -80 °C. Proteins were resolved by 7.5 % or 10 % SDS-PAGE and transferred to 0.2 *μ*m nitrocellulose membranes (Hybond ECL, Amersham Pharmacia Biotech). Membranes were rinsed 3 × with TBS (50 mM Tris-HCl pH 7.6, 145 mM NaCl), blocked with 5 % non-fat dry milk in TBST (TBS / 0.1 % Tween® 20) and probed at 4 °C overnight with the following primary antibodies: mouse monoclonal anti-FLAG M2 (Sigma, 1:1000), mouse monoclonal anti-α-tubulin B512 (Sigma, 1:5000), rat monoclonal anti-tyrosinated α-tubulin YL1/2 (Bio-Rad Laboratories, 1:5000), rabbit polyclonal anti-DYCI-1 (GC1, 1:1000), rabbit polyclonal anti-DNC-1 (GC2, 1:1000), and rabbit polyclonal anti-DNC-2 (GC5, 1:5000). Membranes were washed 5 × with TBST, incubated with goat secondary antibodies coupled to HRP (JacksonImmunoResearch, 1:5000) for 1 hour at room temperature, and washed again 3 × with TBST. Proteins were detected by chemiluminescence using Pierce ECL Western Blotting Substrate (Thermo Scientific) and X-ray film (Fuji).

### Reverse Transcription PCR

Total RNA was isolated from adult hermaphrodites using the TRIzol® Plus RNA Purification Kit (Invitrogen). After 3 washes with M9, pelleted worms were homogenized in 200 *μ*L of TRIzol® reagent with a pellet pestle homogenizer and incubated at room temperature for 5 min. After addition of 40 *μ*L chloroform, samples were shaken vigorously by hand, incubated at room temperature for 3 min, and centrifuged at 12000 x g for 15 min at 4 °C. The upper phase containing the RNA was transferred to an RNase-free tube and an equal volume of 70 % ethanol was added. Further RNA purification steps were performed according to the manufacturer’s instructions. Purified RNA was treated with DNase I (Thermo Scientific), and cDNA was synthesized with the iScript(tm) Select cDNA Synthesis Kit (Bio-Rad Laboratories). The following oligos were used for the PCR reactions in Fig. S2B: forward oligo on *p150*^*dnc-1*^ exon 3 (GAATGTCACCTGCTGCTT); forward oligo on *p150*^*dnc-1*^ exon 4 (AAAGCGGTCTACAACTCC); reverse oligo on *p150*^*dnc-*^*1* exon 5 (GATTGCGATAAGTTGGAGA); reverse oligo on *p150*^*dnc-1*^ exon 6 (AGTAGTCGTGGACGCTTT). For the SL1 PCR shown in Fig. S2D, the following oligos were used: forward oligo on SL1 (GGTTTAATTACCCAAGTTTGA); reverse oligo on *p150*^*dnc-1*^ exon 6 (TCCAGTATCATCAATCTTCTT).

### Embryonic viability

Embryonic viability tests were performed at 20 °C. L4 hermaphrodites were grown on NGM plates with OP50 bacteria for 40 h at 20 °C, then singled-out onto mating plates (NGM plates with a small amount of OP50 bacteria). After 8 h, mothers were removed and the number of hatched and unhatched embryos on each plate was determined 16 h later.

### Live imaging of embryos

Adult gravid hermaphrodite worms were dissected in a watch glass filled with Egg Salts medium (118mM KCl, 3.4 mM MgCl_2_, 3.4 mM CaCl_2_, 5 mM HEPES, pH 7.4), and embryos were mounted onto a 2 % agarose pad. All microscopy was performed in rooms kept at 20 °C. Embryos co-expressing GFP::histone H2B^his-58^ and GFP::γ-tubulin^tbg-1^ were imaged on an Axio Observer microscope (Zeiss) equipped with an Orca Flash 4.0 camera (Hamamatsu), a Colibri.2 light source, and controlled by ZEN software (Zeiss). Embryos expressing GFP::p50^DNC-2^, dynein heavy chain^DHC-1^::GFP, EBP-2::mKate2, and RAB-5::mCherry were imaged on a Nikon Eclipse Ti microscope coupled to an Andor Revolution® XD spinning disk confocal system composed of an iXon Ultra 897 CCD camera (Andor Technology), a solid-state laser combiner (ALC-UVP 350i, Andor Technology), and a CSU-X1 confocal scanner (Yokogawa Electric Corporation), controlled by Andor IQ3 software (Andor Technology).

### Imaging conditions and image analysis

All imaging was performed in one-cell embryos unless otherwise indicated. Image analysis was performed using Fiji software (Image J v1.50b, NIH, Bethesda, Maryland, USA).

#### Pronuclear migration, centrosome positioning, centrosome-centrosome distance, and orientation of centrosome-centrosome axis

Time-lapse sequences of GFP::histone H2B and GFP::γ-tubulin, consisting of 7 × 1 *μ*m z-stacks for GFP fluorescence and one central DIC image captured every 10 s, were recorded at 2 × 2 binning with a 63x oil immersion objective from the start of pronuclear migration until the onset of cytokinesis. Embryo length was defined as the distance between the outermost points of the egg shell visible in the DIC channel. After maximum intensity projection of GFP channel z-stacks, the × and y coordinates of pronuclei and centrosomes were recorded over time using the MTrackJ plugin by manually clicking in the center of the centrosome or nucleus. The position of centrosomes and pronuclei along the anterior-posterior axis was then calculated relative to embryo length, with the anterior reference point set to 0 %. Tracks from individual embryos were aligned relative to pronuclear meeting or nuclear envelope breakdown.

#### Transversal oscillations of the mitotic spindle

Time-lapse sequences of GFP::γ-tubulin, consisting of 12 × 1 *μ*m z-stacks captured every 2 s, were recorded at 2 × 2 binning with a 63x objective from the beginning of metaphase until the end of anaphase. After maximum intensity projection of z-stacks, the × and y coordinates for centrosomes were determined over time with MTrackJ and the transversal distance of each centrosome to a line bisecting the embryo along the anterior-posterior axis was calculated.

#### Levels of GFP::p50^DNC-2^ and dynein heavy chain^DHC-1^::GFP at the nuclear envelope, kinetochores, and on the mitotic spindle

Time-lapse sequences, consisting of 8 × 1 *μ*m z-stacks captured every 10 s, were recorded at 1 × 1 binning with a 60× NA 1.4 oil immersion objective from 40 - 50 s prior to pronuclear meeting until the onset of cytokinesis. Nuclear envelope (NE) signal was quantified 3 frames prior to pronuclear meeting using a maximum intensity projection of the 3 z-sections representing the best in-focus images of the NE. A 2 pixel-wide line was drawn on top of the NE along its entire circumference, and a similar line was drawn next to the NE on the cytoplasmic side around the nucleus. The mean fluorescence signal of the cytoplasmic line was then subtracted from the mean fluorescence signal of the NE line.

Kinetochore signal was measured 7 - 8 frames before the onset of sister chromatid separation using a maximum intensity projection of the z-stack. The top 10 local maxima intensities on kinetochores were identified using the “Find Maxima” function, and the 10 values were averaged. The mean fluorescence intensity of the spindle background close to the kinetochore region was measured and subtracted from the kinetochore signal.

Mitotic spindle signal was measured 2 frames after the onset of chromosome segregation using a maximum intensity projection of the z-stack. The mean intensity in 3 separate 10 × 10 pixel squares on the spindle was determined and the three values were averaged. The mean intensity of three equivalent squares in the cytoplasm adjacent to the spindle served as background signal and was subtracted from the spindle signal.

#### Levels of GFP::p50^DNC-2^, dynein heavy chain^DHC-1^::GFP, and EBP-2::mKate2 at MT plus ends

Time-lapse sequences, consisting of a single cortical confocal section captured every 5 s, were recorded at 1 × 1 binning with a 100× NA 1.4 oil immersion objective for 1 min beginning at metaphase. 3 different images at least 3 frames apart from each other were used for quantifications. A circle with a 5-pixel radius was drawn around individual MT plus ends (marked by EBP-2::mKate2) and the integrated fluorescence intensity was measured in each channel. The circle was then expanded by increasing the radius by 2 pixels, and the integrated intensity of this larger circle was measured in each channel. Background was defined as the difference in integrated intensities between the larger and the smaller circle. The background value was scaled in proportion to the smaller circle and then subtracted from the integrated intensity of the smaller circle to obtain a final value for both channels. For each MT plus end, the GFP value was then divided by the mKate2 value, except for the p150^DNC-1^ isoform mutants (Fig. 2D) and the α-tubulin tyrosine mutant (Fig. 4I), where only the GFP channel was used for measurements.

#### Cortical residency times of GFP::p50^DNC-2^, dynein heavy chain^DHC-1^::GFP, and EBP- 2::mKate2

Time-lapse sequences, consisting of a single cortical confocal section captured every 200 ms, were recorded at 1 × 1 binning with a 100× NA 1.4 oil immersion objective for 1 min beginning at metaphase. Image sequences were analyzed using the LoG detector (estimated blob diameter 10 pixels) and the Simple LAP Tracker (linking max distance 5 pixels; gap-closing max distance 5 pixels; gap-close max frame gap 0 frames) in the TrackMate plugin. The values for track duration were defined as the cortical residency times for GFP and mKate2 puncta.

#### Levels of GFP::p50^DNC-2^ and dynein heavy chain^DHC-1^::GFP at the EMS-P2 cell border

Time-lapse sequences, consisting of 8 × 1 *μ*m z-stacks captured every 30 s, were recorded at 1 × 1 binning with 60× NA 1.4 oil immersion objective. Four-cell embryos were imaged from the beginning of nuclear envelope breakdown in AB cells until cytokinesis of the P2 cell. The signal at the EMS-P2 cell border was measured at the time of EMS spindle rotation in maximum intensity projections of the z-stacks. The top 10 local maxima intensities on the EMS-P2 cell border were identified using the “Find Maxima” function in Fiji, and the 10 values were averaged. The mean fluorescence intensity of an adjacent area (20 × 20 pixels) in the EMS cytoplasm served as background and was subtracted from the EMS-P2 cell border signal.

#### Tracking of early endosomes marked with mCherry::RAB-5

Time-lapse sequences, consisting of a single confocal section captured every 400 ms, were recorded at 1 × 1 binning with a 60× NA 1.4 oil immersion objective for 6 min beginning at pronuclear meeting. Image sequences were analyzed using the LoG detector (estimated blob diameter 6 pixels) and the Simple LAP Tracker (linking max distance 8 pixels; gap-closing max distance 16 pixels; gap-close max frame gap 2 frames) in the TrackMate plugin. All tracks whose particles showed directed movement towards centrosomes and had a track displacement of at least 0.9 *μ*m (5 pixels) were considered.

### Statistical analysis

Values in figures and text are reported as mean ± SEM with a 95 % confidence interval. Statistical significance was evaluated with the t-test using GraphPad Prism 7 software.

## ACKNOWLEDGEMENTS

The research leading to these results has received funding from the European Research Council under the European Union’s Seventh Framework Programme (FP7/2007-2013)/ ERC grant agreement n° ERC-2013-StG-338410-DYNEINOME. Additional funding was provided by an EMBO Installation Grant (R.G.) and by the Fundação para a Ciência e a Tecnologia to R. G. (IF/01015/2013/CP1157/CT0006) and D. J. B. (SFRH/BPD/101898/2014). Some *C. elegans* strains were provided by the CGC, which is funded by NIH Office of Research Infrastructure Programs (P40 OD010440).

The authors declare no competing financial interests.

## SUPPLEMENTAL FIGURE LEGENDS

**Figure S1:**
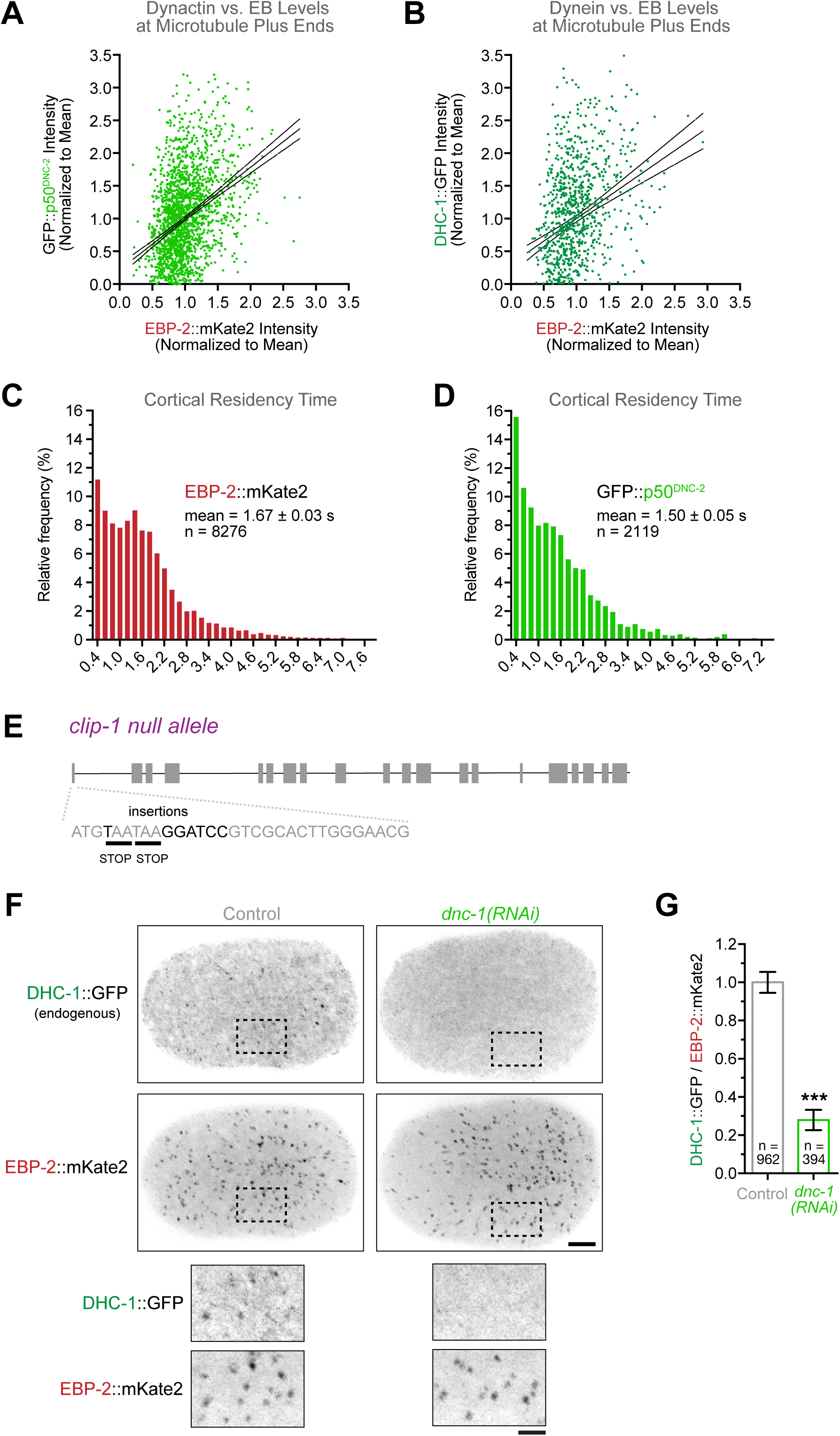
Dynein targeting to MT plus ends depends on dynactin. **(A), (B)** Correlation plots of GFP::p50^DNC-2^ versus EBP-2::mKate2 intensity *(A)* and Dynein Heavy Chain^DHC-1^::GFP versus EBP-2::mKate2 intensity *(B)*, measured at the cortex of metaphase one-cell embryos. **(C), (D)** Residency time of EBP-2::mKate2 *(C)* and GFP::p50^DNC-2^ *(D)* puncta at the cortex of metaphase one-cell embryos. The total number *(n)* of plus ends scored is indicated, derived from at least 8 embryos. **(E)** Schematic of the *clip-1* locus. The mutations introduced by CRISPR-Cas9-based genome editing to generate a null allele are indicated in black font. **(F)** Cortical confocal section of one-cell embryos in metaphase co-expressing endogenous dynein heavy chain^DHC-1^::GFP and EBP-2::mKate2, showing that depletion of p150^DNC-1^ delocalizes dynein from MT tips. Image is a maximum intensity projection over time of 12 images acquired every 5 s. Scale bar, 5 *μ*m; insets, 2 *μ*m. **(G)** Quantification of dynein heavy chain^DHC-1^::GFP levels at MT plus ends using fluorescence intensity measurements at the cortex. Error bars represent the SEM with a 95 % confidence interval, and *n* indicates the total number of MT plus ends measured in 7-8 embryos per condition. The t-test was used to determine statistical significance (*** indicates p < 0.0001).

**Figure S2:**
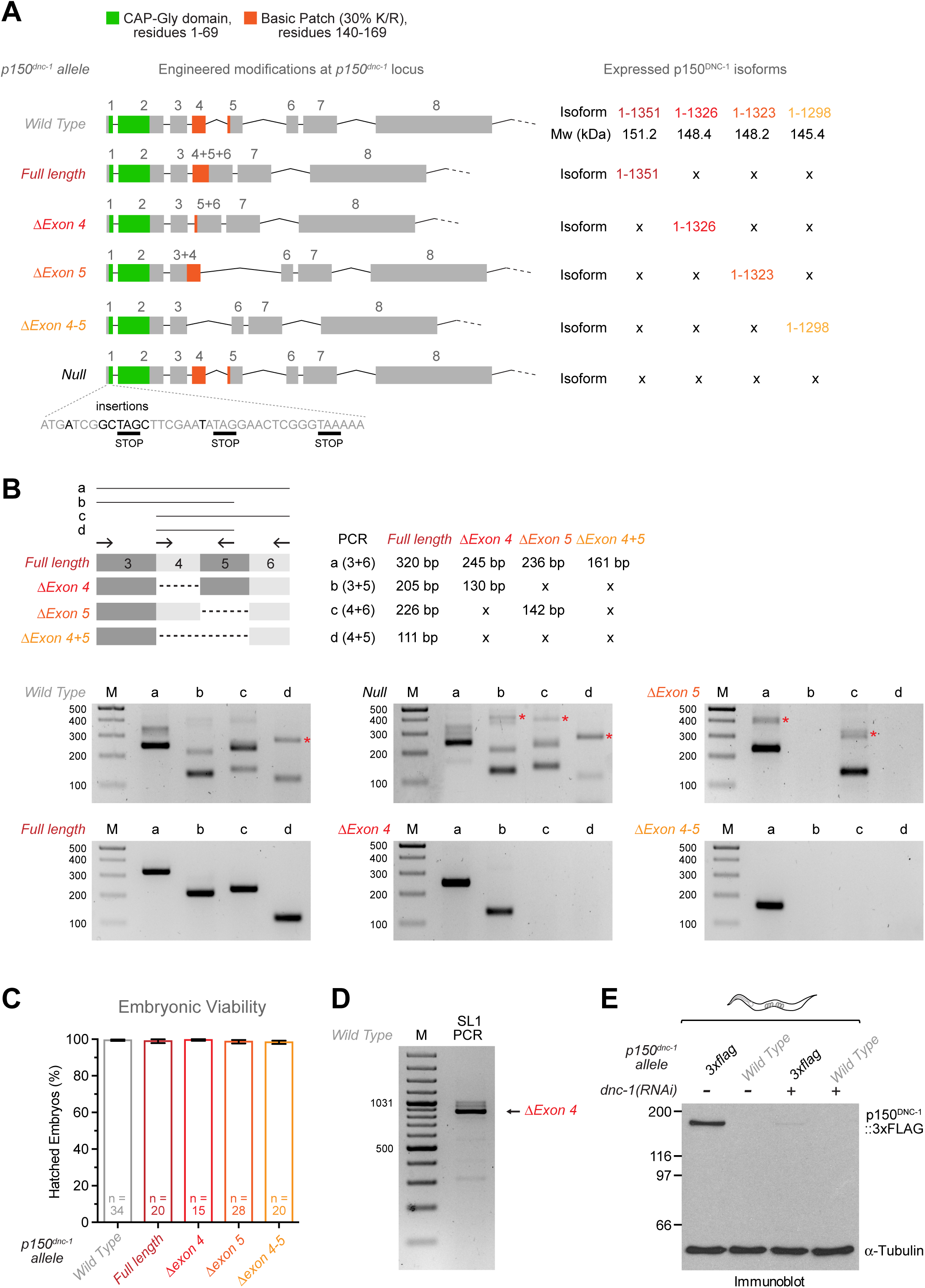
Engineering of mutants that restrict p150^DNC-1^ expression to single splice isoforms. **(A)** Schematic of the *p150*^*dnc-1*^ locus with engineered modifications. By deleting and/or fusing exons, p150^DNC-1^ expression was restricted to single N-terminal splice isoforms (full length, Δexon 4, Δexon 5, or Δexons4-5), as indicated on the right. In addition, introduction of a frameshift mutation after the *p150*^*dnc-1*^ start codon generated a null allele. **(B)** Results of reverse transcription PCRs using RNA isolated from adult worms and primer pairs that allow detection of the four splice isoforms. Primer locations and predicted sizes of PCR products for the different isoforms are indicated. Crosses (*x*) indicate that the PCR will not amplify any product. All four N-terminal splice isoforms are detected in wild-type and *p150*^*dnc-1*^ null mutant adults, whereas only one isoform is detected in each *p150*^*dnc-1*^ isoform mutant. Asterisks next to gel bands denote unspecific PCR products. M, DNA size marker. **(C)** Embryonic viability assay for *p150*^*dnc-1*^ isoform mutants. Error bars represent the SEM with a 95 % confidence interval, and *n* indicates the number of hermaphrodite mothers whose progeny was counted for each condition (> 500 total progeny per condition). **(D)** Result of a reverse transcription PCR using RNA isolated from adult wild-type worms with one of the primers recognizing the spliced leader sequence 1 (SL1) and the other located in exon 6 of *p150*^*dnc-1*^. In *C. elegans*, about 70 % of mRNAs are trans-spliced to one of two 22 nucleotide spliced leaders, SL1 or SL2, which replace the 5’ ends of pre-mRNAs. One major product was amplified in the SL1 PCR and identified as the Δ*exon4* isoform by DNA sequencing. A PCR reaction with a primer recognizing SL2 did not amplify any product (data not shown). M, DNA size marker. **(E)** Immunoblot of wild-type or *p150*^*dnc-1*^*::3xflag* adult worms with an antibody against the 3xFLAG tag, showing that there is no detectable p150^DNC-1^ isoform corresponding to human p135. α-Tubulin was used as the loading control. Molecular mass is indicated in kilodaltons.

**Figure S3:**
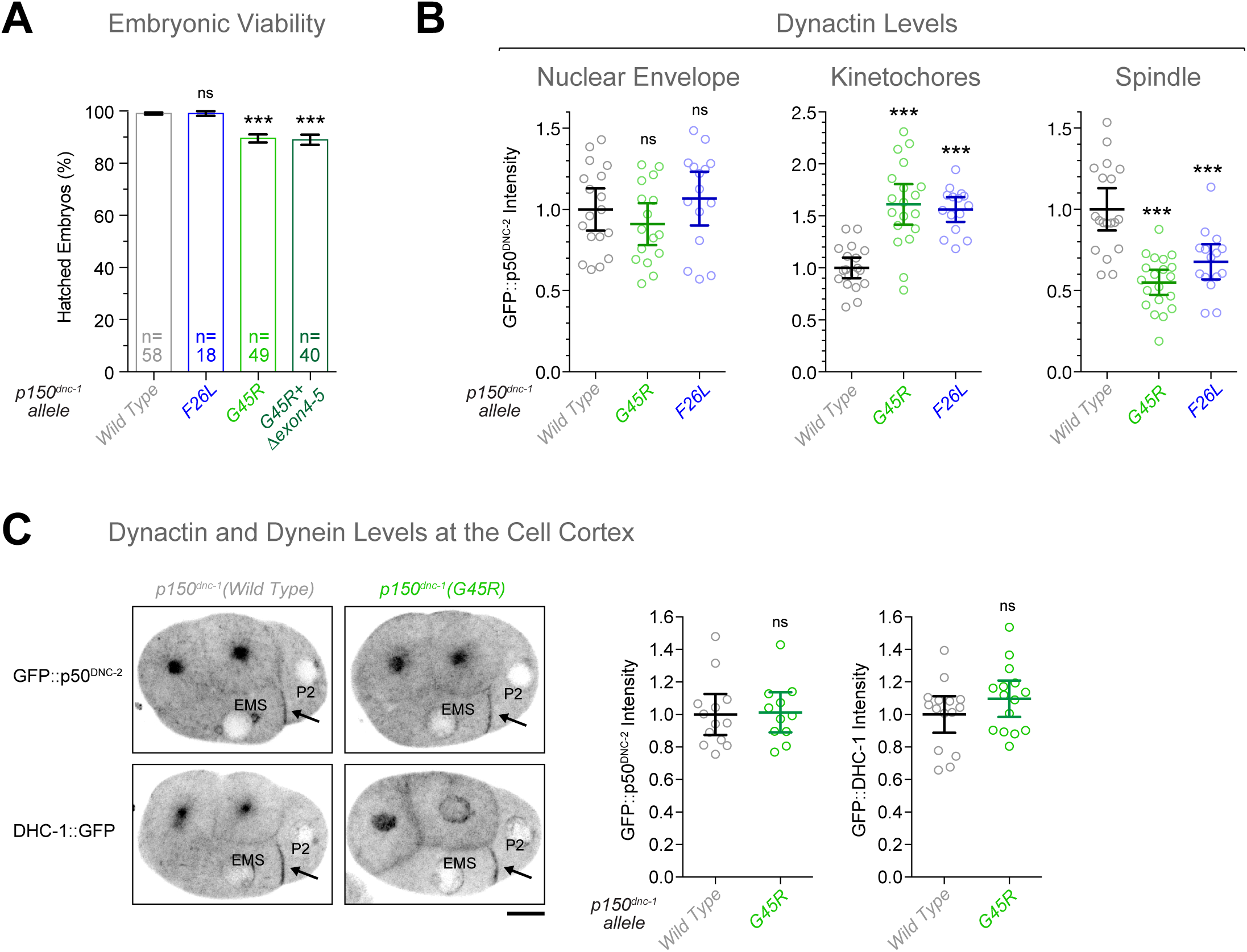
MT tip tracking of p150^DNC-1^ is dispensable for targeting dynactin and dynein to the cell cortex. **(A)** Embryonic viability assay for p150^DNC-1^ CAP-Gly domain mutants. Error bars represent the SEM with a 95 % confidence interval, and *n* indicates the number of hermaphrodite mothers whose progeny was counted for each condition (> 500 total progeny per condition). The t-test was used to determine statistical significance (*** indicates p < 0.0001; ns = not significant, p > 0.05). **(B)** Quantification of dynactin levels at the nuclear envelope, kinetochores, and the mitotic spindle for the p150^DNC-1^ mutants G45R and F26L, using fluorescence intensity measurements of GFP::p150^DNC-2^. Circles represent measurements in individual embryos. Error bars represent the SEM with a 95 % confidence interval. The t-test was used to determine statistical significance (*** indicates p < 0.0001; ns = not significant, p > 0.05). ***(C)*** *(left)* Stills from a time-lapse sequence in 4-cell embryos expressing GFP::p50^DNC-2^ *(top)* or dynein heavy chain^DHC-1^::GFP *(bottom)*, showing normal accumulation of dynein-dynactin at the EMS-P2 cell border in the p150^dnc-^ 1(G45R) mutant. Scale bar, 5 *μ*m. *(right)* Quantification of dynein and dynactin levels at the EMS-P2 cell border using fluorescence intensity measurements. Error bars represent the SEM with a 95 % confidence interval. The t-test was used to determine statistical significance (ns = not significant, p > 0.05).

**Figure S4:**
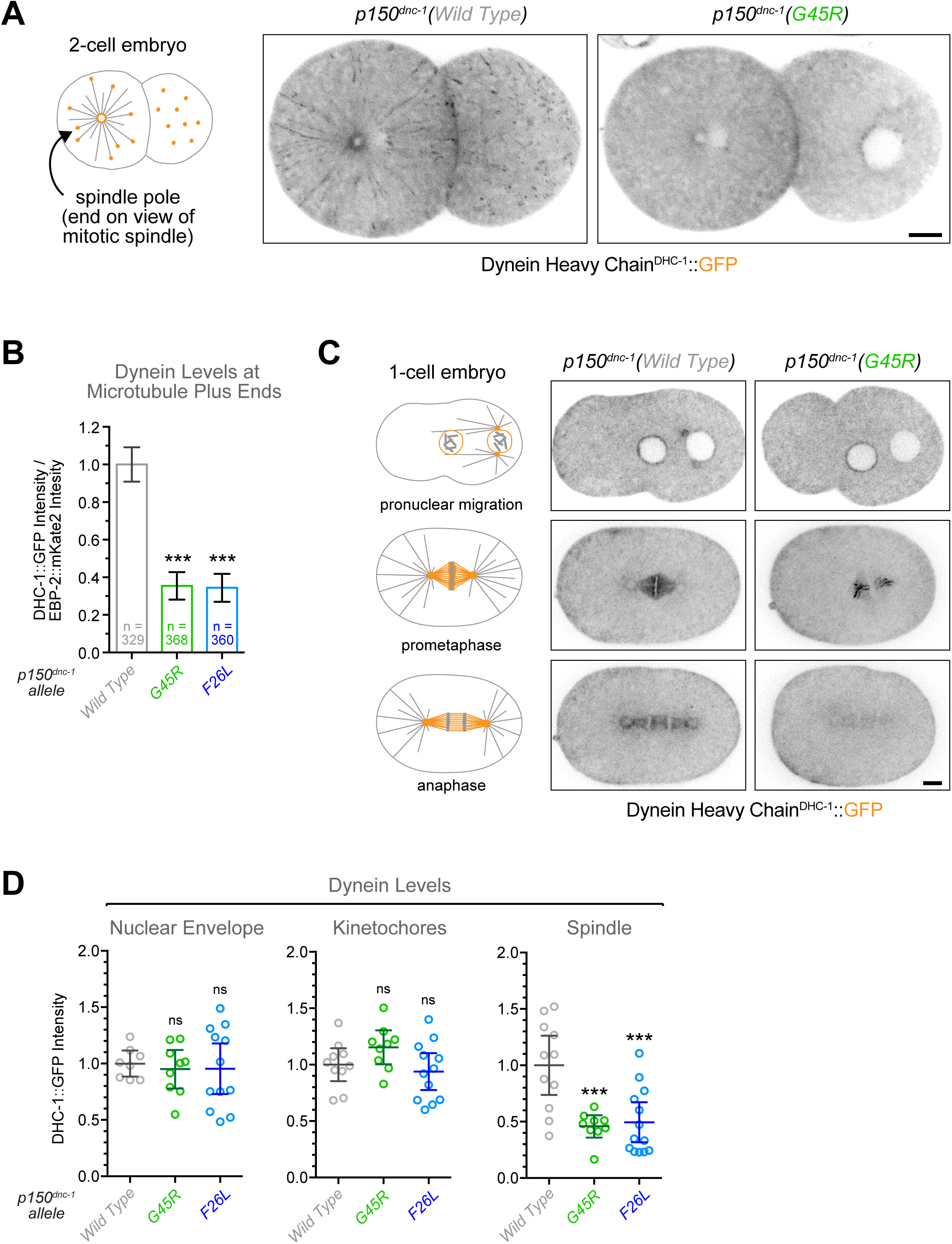
p150^DNC-1^ CAP-Gly domain mutants delocalize dynein from MT tips. **(A)** Stills from a time-lapse sequence in 2-cell embryos expressing dynein heavy chain^DHC-1^::GFP, demonstrating that dynein is delocalized from MT plus ends in the p150^DNC-1^ G45R mutant. Scale bar, 5 *μ*m. **(B)** Quantification of dynein levels at MT plus ends using fluorescence intensity measurements of dynein heavy chain^DHC-1^::GFP at the cortex of metaphase one-cell embryos. For each MT end, dynein heavy chain^DHC-1^::GFP signal was normalized to EBP-2::mKate2 signal. Error bars represent the SEM with a 95 % confidence interval, and *n* indicates the number of individual MT ends analyzed from 6-7 embryos per condition. The t-test was used to determine statistical significance (*** indicates p < 0.0001). **(C)** Stills from time-lapse sequences in one-cell embryos expressing dynein heavy chain^DHC-1^::GFP, showing that the p150^DNC-1^ G45R mutant reduces dynein levels on the mitotic spindle and at centrosomes, but not at the nuclear envelope and kinetochores. Scale bar, 5 *μ*m. **(D)** Quantification of dynein levels at the nuclear envelope, kinetochores, and the mitotic spindle for the p150^DNC-1^ mutants G45R and F26L, using fluorescence intensity measurements for dynein heavy chain^DHC-1^::GFP in images as shown in *(C)*. Circles represent measurements in individual embryos. Error bars represent the SEM with a 95 % confidence interval. The t-test was used to determine statistical significance (*** indicates p < 0.0001; ns = not significant, p > 0.05).

**Figure S5:**
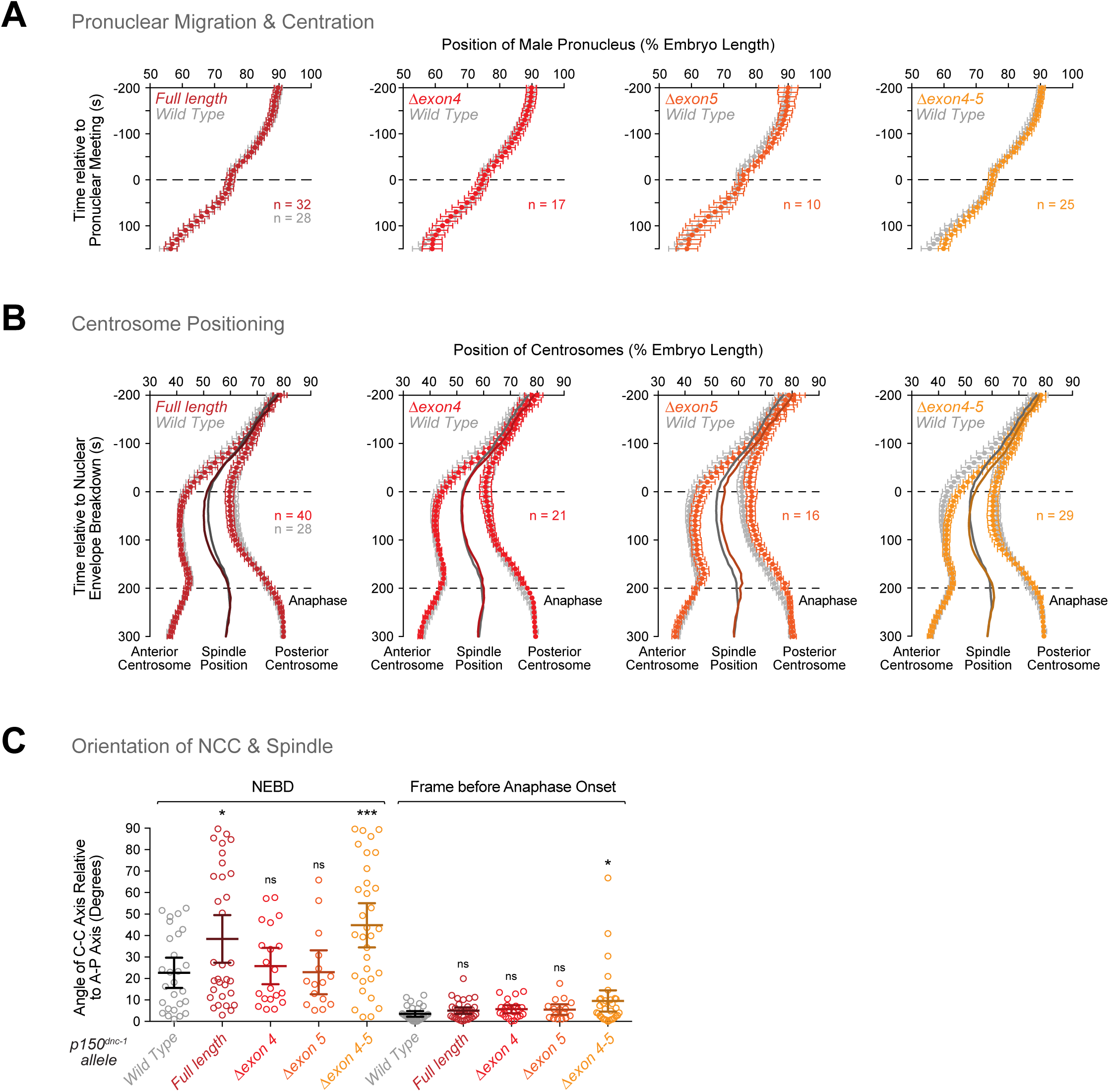
Functional analysis of p150^DNC-1^ isoform mutants. **(A)** Migration kinetics of the male pronucleus in one-cell embryos expressing single isoforms of p150^DNC-1^. The position of the male pronucleus, marked by GFP::histone H2B, was determined along the anterior-posterior axis in images captured every 10 s. Individual traces were normalized to embryo length, time-aligned relative to pronuclear meeting, averaged for the indicated number (n) of embryos, and plotted against time. Error bars represent the SEM with a 95 % confidence interval. **(B)** Positioning of centrosomes, marked by GFP::γ-tubulin, measured in time-lapse sequences as described for *(A)* and plotted relative to NEBD. Solid lines indicate the midpoint between the two centrosomes (spindle position). Anaphase begins at 200 s. Error bars represent the SEM with a 95 % confidence interval. **(C)** Angle between the centrosome-centrosome (C-C) axis and the anterior-posterior (A-P) axis in one-cell embryos at NEBD and anaphase onset. Circles correspond to measurements in individual embryos. Error bars represent the SEM with a 95 % confidence interval. The t-test was used to determine statistical significance (*** indicates p < 0.0001; * indicates p < 0.05; ns = not significant, p > 0.05).

**Figure S6:**
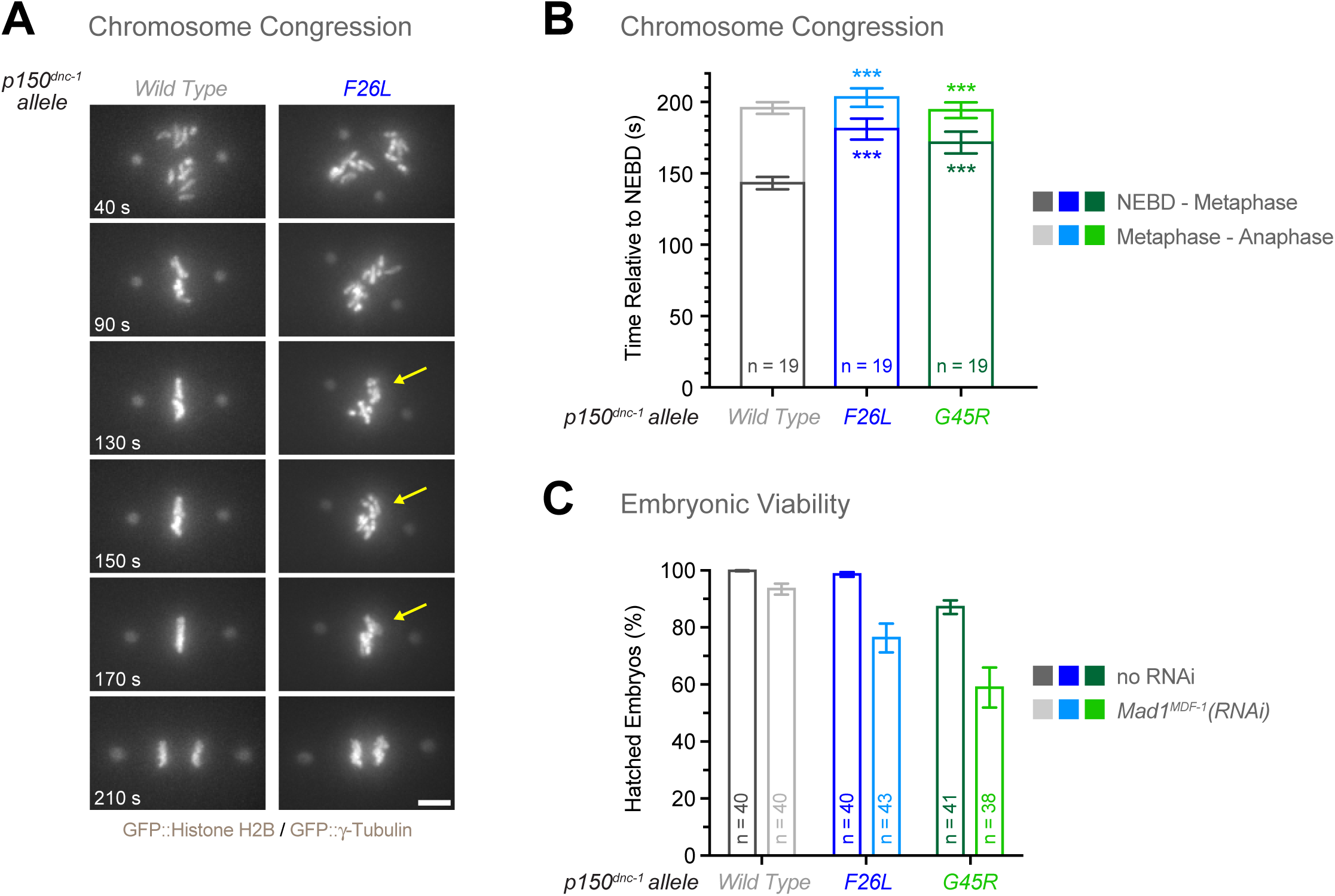
p150^DNC-1^ CAP-Gly mutants delay chromosome congression. **(A)** Selected frames from time-lapse sequences in one-cell embryos expressing GFP::histone H2B and GFP::γ-tubulin, showing that congression of chromosomes is delayed in the p150^DNC-1^ F26L mutant. Time is relative to NEBD. Scale bar, 5 *μ*m. **(B)** Interval duration for NEBD to metaphase (full alignment of chromosomes) and metaphase to anaphase (onset of sister chromatid separation). Error bars represent the SEM with a 95 % confidence interval, and *n* indicates the number of embryos analyzed. **(C)** Embryonic viability assay for p150^DNC-1^ CAP-Gly mutants with and without depletion of the SAC component Mad1^MDF-1^. Error bars represent the SEM with a 95 % confidence interval, and *n* indicates the number of hermaphrodite mothers whose progeny was counted for each condition (> 500 total progeny per condition).

**Figure S7:**
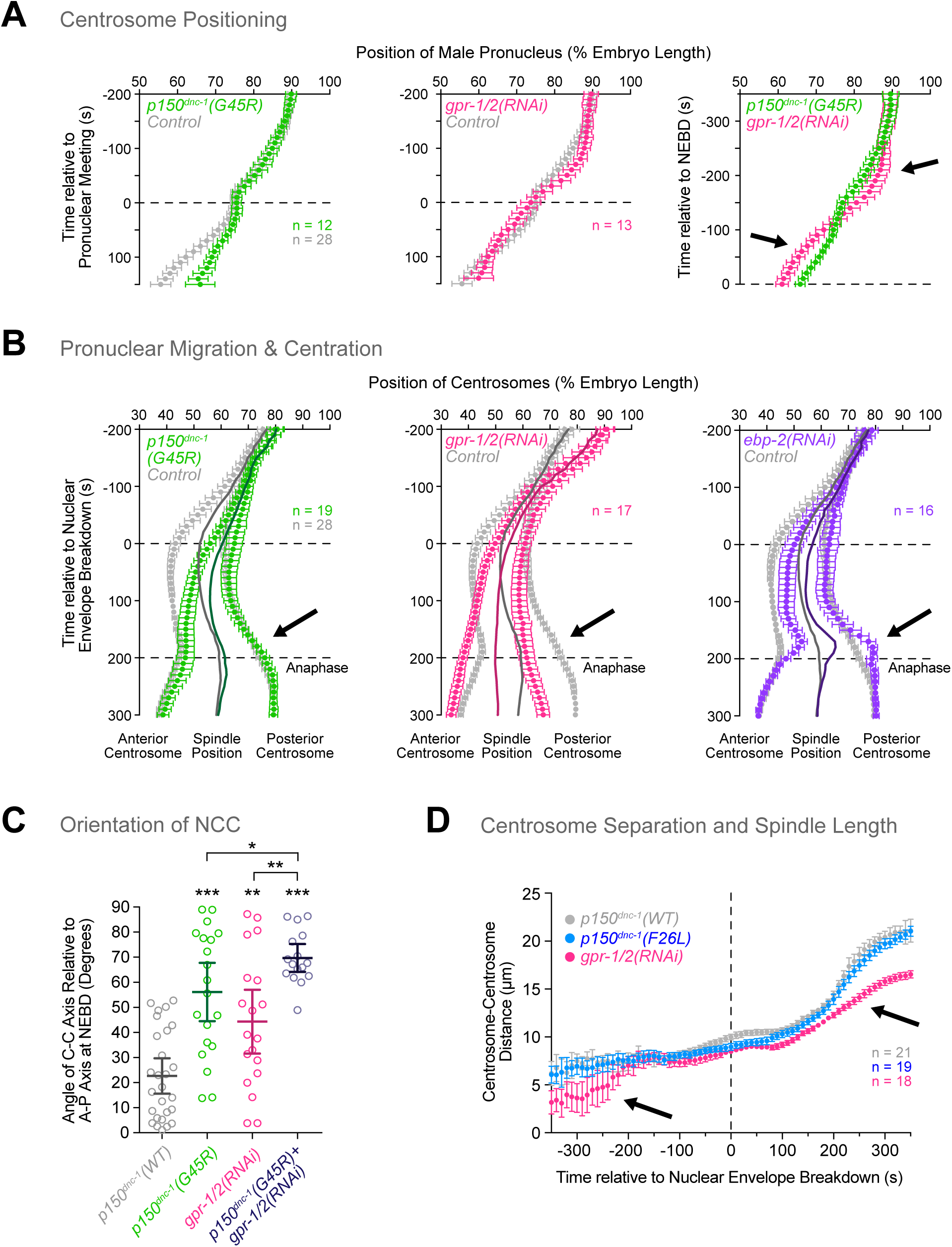
Inhibition of cortical pulling forces and p150^DNC-1^ CAP-Gly mutants cause distinct defects. **(A)** Migration kinetics of the male pronucleus in one-cell embryos. The position of the male pronucleus, marked by GFP::histone H2B, was determined along the anterior-posterior axis in images captured every 10 s. Individual traces were normalized to embryo length, time-aligned relative to pronuclear meeting *(left and middle)* or NEBD *(right)*, averaged for the indicated number (n) of embryos, and plotted against time. Arrows point to differences in migration kinetics between the p150^DNC-1^ G45R mutant and *gpr-1/2(RNAi)*. Error bars represent the SEM with a 95 % confidence interval. **(B)** Positioning of centrosomes, marked by GFP::γ-tubulin, measured in time-lapse sequences as described for *(A)* and plotted relative to NEBD. Solid lines indicate the midpoint between the two centrosomes (spindle position). Error bars represent the SEM with a 95 % confidence interval. Arrows highlight the difference in posterior centrosome displacement between the p150^DNC-1^ G45R mutant, *gpr-1/2(RNAi)*, and *ebp-2(RNAi)*. **(C)** Angle between the centrosome-centrosome (C-C) axis and the anterior-posterior (A-P) axis in one-cell embryos at NEBD. Circles correspond to measurements in individual embryos. Error bars represent the SEM with a 95 % confidence interval. The t-test was used to determine statistical significance (*** indicates p < 0.0001; ** indicates p < 0.01; * indicates p < 0.05). **(D)** Plot of centrosome-centrosome distance over time in one-cell embryos expressing GFP::γ-tubulin. Measurements were made in images captured every 10 s. Individual traces were time-aligned relative to NEBD, averaged for the indicated number (n) of embryos, and plotted against time. Error bars represent the SEM with a 95 % confidence interval. Arrows points to delays in centrosome separation (-300 s) and mitotic spindle elongation (200 s) in embryos depleted of GPR-1/2.

**Movie S1: Microtubule tip tracking of dynactin in the *C. elegans* embryo.**

One-cell embryo in metaphase co-expressing GFP::p50^DNC-2^ and EBP-2::mKate2. The anterior side of the embryo is to the left. A single confocal section in the embryo center was acquired every 0.4 s. Playback speed is 30 frames per second.

**Movie S2: MT tip tracking of dynein in the early *C. elegans* embryo.**

Two-cell embryo expressing dynein heavy chain^DHC-1^::GFP. The anterior side of the embryo is to the left. A single confocal section near the cortex was acquired every 0.2 s. Playback speed is 30 frames per second.

**Movie S3: p150^DNC-1^ CAP-Gly mutants delocalize dynactin from microtubule tips - center view.**

One-cell embryos in metaphase expressing GFP::p50^DNC-2^. The anterior side of the embryo is to the left. A single confocal section in the embryo center was acquired every 0.2 s. Playback speed is 10 frames per second.

**Movie S4: p150^DNC-1^ CAP-Gly mutants delocalize dynactin from microtubule tips - cortical view.**

One-cell embryos in metaphase expressing GFP::p50^DNC-2^. The anterior side of the embryo is to the left. A single confocal section at the cortex was acquired every 0.2 s. Playback speed is 10 frames per second.

**Movie S5: Centration/rotation defects in p150^DNC-1^ CAP-Gly mutants.**

Mitotic one-cell embryos expressing GFP-labelled histone H2B and γ-Tubulin to mark chromosomes and centrosomes, respectively. The anterior side of the embryos is to the left. Time lapse is 10 s and the playback speed is 6 frames per second.

**Movie S6: Delayed chromosome congression and dampened anaphase spindle rocking in p150^DNC-1^ CAP-Gly mutants.**

Mitotic one-cell embryos expressing GFP-labelled histone H2B and γ-Tubulin to mark chromosomes and centrosomes, respectively. The anterior side of the embryos is to the left. Time lapse is 2 s and the playback speed is 6 frames per second.

**Movie S7: Centration/rotation defects in α-tubulin tyrosine mutants.**

Mitotic one-cell embryos expressing GFP-labelled histone H2B and γ-Tubulin to mark chromosomes and centrosomes, respectively. The anterior side of the embryos is to the left. Time lapse is 10 s and the playback speed is 6 frames per second.

**Movie S8: Defective transport of early endosomes in p150^DNC-1^ CAP-Gly mutants.**

One-cell embryos during the centration phase expressing mCherry::RAB-5 to label early endosomes. The anterior side of the embryos is to the left. Time lapse is 0.4 s and playback speed is 50 frames per second.

**Supplemental Table S1:** worm strains used in this study

| Strain name | Genotype |
| --- | --- |
| N2 | WT (ancestral N2 Bristol) |
| GCP181 | dnc-1[prt7(F26L)] IV |
| GCP231 | dnc-1[prt12(G33S)] IV; unc-5(e53) IV/nT1 [qls51] (IV;V); dpy-11(e224) V/nT1 [qls51] (IV;V) |
| GCP237 | dnc-1[prt14(Δexons 4-5)] IV |
| GCP242 | dnc-1[prt15(C-terminal 3xflag)] IV |
| GCP245 | dnc-1[prt18(G45R)] IV |
| GCP247 | dnc-1[prt8(exons 5-6 fusion)] IV; dnc-1[prt20(exons 4-5 fusion)] IV |
| GCP255 | dnc-1[prt23(knockout)]; unc-5(e53) IV/nT1 [qls51] (IV;V); dpy-11(e224) V/nT1 [qls51] (IV;V) |
| GCP289 | dnc-1[prt7(F26L)] IV; unc-119(ed3) III; ruls32[pAZ132; Ppie-1::gfp::his-58; unc-119(+)] III; ddls6 [Ppie-1::gfp::tbG-1; unc-119(+)] V |
| GCP290 | dnc-1[prt18(G45R)] IV; unc-119(ed3) III; ruls32[pAZ132; Ppie-1::gfp::his-58; unc-119(+)] III; ddls6 [Ppie-1::gfp::tbG-1; unc-119(+)] V |
| GCP291 | dnc-1[prt14(Δexons 4-5)] IV; unc-119(ed3) III; ruls32[pAZ132; Ppie-1::gfp::his-58; unc-119(+)] III; ddls6 [Ppie-1::gfp::tbG-1; unc-119(+)] V |
| GCP292 | dnc-1[prt8(exons 5-6 fusion)] IV; dnc-1[prt20(exons 4-5 fusion)] IV; unc-119(ed3) III; ruls32[pAZ132; Ppie-1::gfp::his-58; unc-119(+)] III; ddls6 [Ppie-1::gfp::tbG-1; unc-119(+)] V |
| GCP412 | dnc-1[prt7(F26L)] IV; dhc-1[C-terminal gfp] I |
| GCP413 | dnc-1[prt18(G45R)] IV; dhc-1[C-terminal gfp] I |
| GCP416 | unc-119(ed3) III; prtSi122[pRG629; Pmex-5::ebp-2::mkate2::tbb-2 3'UTR; cb-unc-119(+)] II |
| GCP417 | dnc-2[prt42(N-terminal 3xflag::gfp)] III |
| GCP430 | dnc-1[prt7(F26L)] IV; unc-119(ed3) III; prtSi122[pRG629; Pmex-5::ebp-2::mkate2::tbb-2 3'UTR; cb-unc-119(+)] II |
| GCP431 | dnc-1[prt18(G45R)] IV; unc-119(ed3) III; prtSi122[pRG629; Pmex-5::ebp-2::mkate2::tbb-2 3'UTR; cb-unc-119(+)] II |
| GCP432 | dnc-1[prt7(F26L)] IV; dnc-2[prt42(N-terminal 3xflag::gfp)] III |
| GCP433 | dnc-1[prt18(G45R)] IV; dnc-2[prt42(N-terminal 3xflag::gfp)] III |
| GCP442 | tba-1[prt55(Y454A)] I |
| GCP444 | dnc-1[prt14(Δexons 4-5)] IV; dnc-2[prt42(N-terminal 3xflag::gfp)] III |
| GCP445 | dnc-1[prt8(exons 5-6 fusion)] IV; dnc-1[prt20(exons 4-5 fusion)] IV; dnc-2[prt42(N-terminal 3xflag::gfp)] III |
| GCP446 | dhc-1[C-terminal gfp] I; unc-119(ed3) III; prtSi122[pRG629; Pmex-5::ebp-2::mkate2::tbb-2 3'UTR; cb-unc-119(+)] II |
| GCP447 | dnc-2[prt42(N-terminal 3xflag::gfp)] III; unc-119(ed3) III; prtSi122[pRG629; Pmex-5::ebp-2::mkate2::tbb-2 3'UTR; cb-unc-119(+)] II |
| GCP452 | tba-2[prt57(Y448A)] I |
| GCP477 | dnc-1[prt8(exons 5-6 fusion)] IV; dnc-1[prt61(Δexon 4)] IV |
| GCP487 | tba-1[prt55(Y454A)] I; tba-2[prt58(Y448A)] I |
| GCP494 | dnc-1[prt69(exons 3-4 fusion)] IV; dnc-1[prt71(Δexon 5)] IV |
| GCP499 | dnc-1[prt8(exons 5-6 fusion)] IV; dnc-1[prt61(Δexon 4)] IV; dnc-2[prt42(N-terminal 3xflag::gfp)] III |
| GCP500 | dnc-1[prt8(exons 5-6 fusion)] IV; dnc-1[prt61(Δexon 4)] IV; unc-119(ed3) III; ruls32[pAZ132; Ppie-1::gfp::his-58; unc-119(+)] III; ddls6 [Ppie-1::gfp::tbG-1; unc-119(+)] V |
| GCP502 | tba-1[prt55(Y454A)] I; tba-2[prt58(Y448A)] I; dnc-2[prt42(N-terminal 3xflag::gfp)] III |
| GCP503 | tba-1[prt55(Y454A)] I; tba-2[prt58(Y448A)] I; unc-119(ed3) III; ruls32[pAZ132; Ppie-1::gfp::his-58; unc-119(+)] III; ddls6 [Ppie-1::gfp::tbG-1; unc-119(+)] V |
| GCP520 | dnc-1[prt69(exons 3-4 fusion)] IV; dnc-1[prt71(Δexon 5)] IV; dnc-2[prt42(N-terminal 3xflag::gfp)] III |
| GCP522 | dnc-1[prt69(exons 3-4 fusion)] IV; dnc-1[prt71(Δexon 5)] IV; unc-119(ed3) III; ruls32[pAZ132; Ppie-1::gfp::his-58; unc-119(+)] III; ddls6 [Ppie-1::gfp::tbG-1; unc-119(+)] V |
| GCP534 | dnc-1[prt7(F26L)] IV; dnc-2[prt42(N-terminal 3xflag::gfp)] III; unc-119(ed3) III; prtSi122[pRG629; Pmex-5::ebp-2::mkate2::tbb-2 3'UTR; cb-unc-119(+)] II |
| GCP535 | dnc-1[prt18(G45R)] IV; dnc-2[prt42(N-terminal 3xflag::gfp)] III; unc-119(ed3) III; prtSi122[pRG629; Pmex-5::ebp-2::mkate2::tbb-2 3'UTR; cb-unc-119(+)] II |
| GCP562 | dnc-1[prt18(G45R)] IV; dnc-1[prt74(Δexons 4-5)] IV; unc-119(ed3) III; ruls32[pAZ132; Ppie-1::gfp::his-58; unc-119(+)] III; ddls6 [Ppie-1::gfp::tbG-1; unc-119(+)] V |
| GCP575 | dnc-1[prt18(G45R)] IV; tba-1[prt55(Y454A)] I; tba-2[prt58(Y448A)] I; unc-119(ed3) III; ruls32[pAZ132; Ppie-1::gfp::his-58; unc-119(+)] III; ddls6 [Ppie-1::gfp::tbG-1; unc-119(+)] V |
| GCP581 | dnc-1[prt7(F26L)] IV; dhc-1[C-terminal gfp] I; unc-119(ed3) III; prtSi122[pRG629; Pmex-5::ebp-2::mkate2::tbb-2 3'UTR; cb-unc-119(+)] II |
| GCP582 | dnc-1[prt18(G45R)] IV; dhc-1[C-terminal gfp] I; unc-119(ed3) III; prtSi122[pRG629; Pmex-5::ebp-2::mkate2::tbb-2 3'UTR; cb-unc-119(+)] II |
| GCP585 | dnc-1[prt18(G45R)] IV; dnc-1[prt74(Δexons 4-5)] IV; dnc-2[prt42(N-terminal 3xflag::gfp)] III; unc-119(ed3) III; prtSi122[pRG629; Pmex-5::ebp-2::mkate2::tbb-2 3'UTR; cb-unc-119(+)] II |
| GCP586 | dnc-1[prt18(G45R)] IV; dnc-1[prt74(Δexons 4-5)] IV; unc-119(ed3) III; ltIs79[pAA196; Ppie-1::mcherry::rab-5; unc-119(+)]; pwIs116[Prme-2::rme-2::gfp::rme-2 3'UTR; unc-119(+)] |
| GCP587 | tba-1[prt55(Y454A)] I; tba-2[prt58(Y448A)] I; unc-119(ed3) III; ltIs79[pAA196; Ppie-1::mcherry::rab-5; unc-119(+)]; pwIs116[Prme-2::rme-2::gfp::rme-2 3'UTR; unc-119(+)] |
| GCP602 | clip-1[prt92(knockout)] III; dnc-2[prt42(N-terminal 3xflag::gfp)] III; unc-119(ed3) III; prtSi122[pRG629; Pmex-5::ebp-2::mkate2::tbb-2 3'UTR; cb-unc-119(+)] II |
| OD179 | unc-119(ed3) III; ltIs79[pAA196; Ppie-1::mcherry::rab-5; unc-119(+)]; pwIs116[Prme-2::rme-2::gfp::rme-2 3'UTR; unc-119(+)] |
| OD2955 | dhc-1[C-terminal gfp] I |
| TH32 | unc-119(ed3) III; ruls32[pAZ132; Ppie-1::gfp::his-58; unc-119(+)] III; ddls6[Ppie-1::gfp::tbG-1; unc-119(+)] V |

**Supplemental Table S2:**
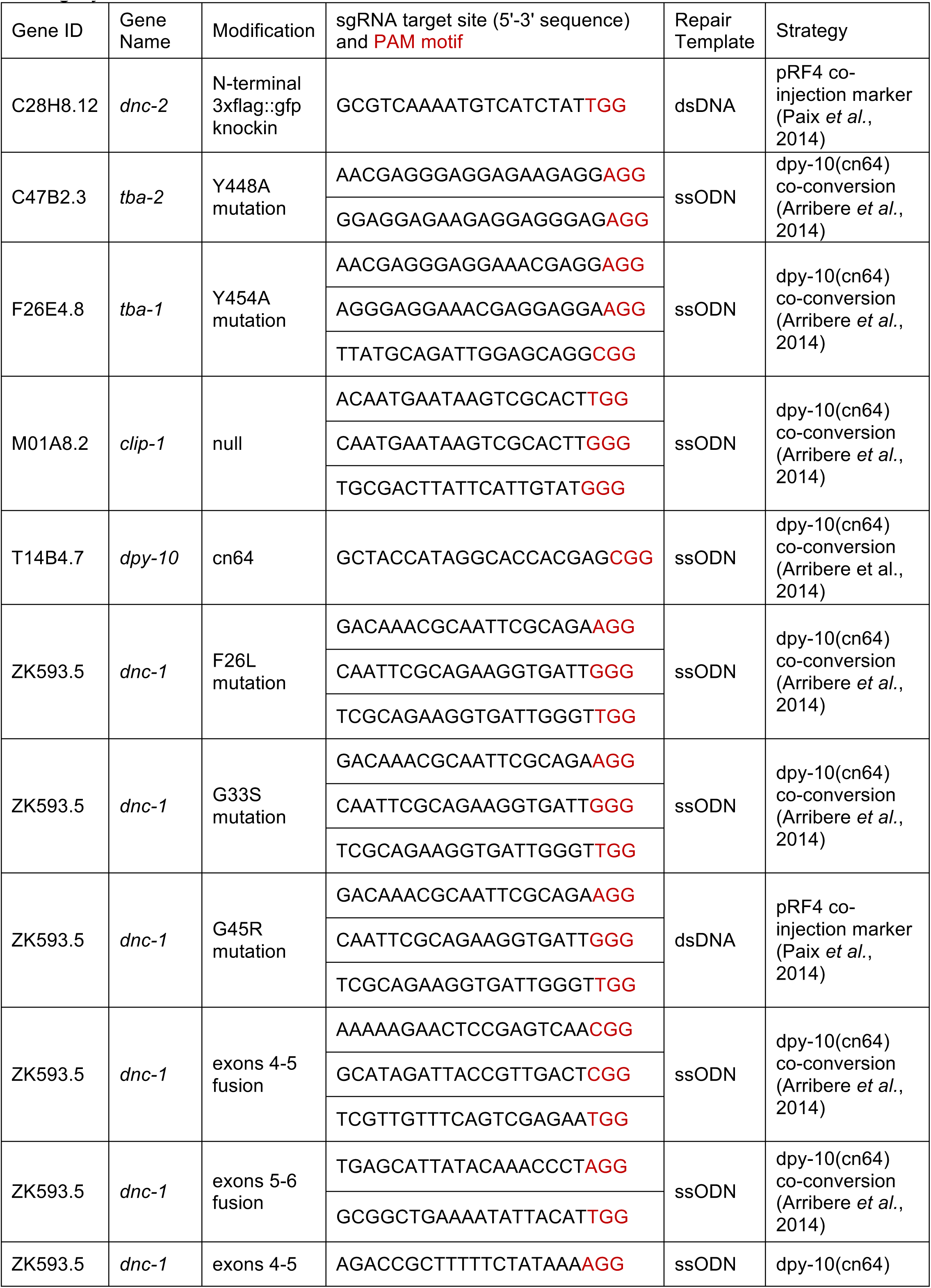
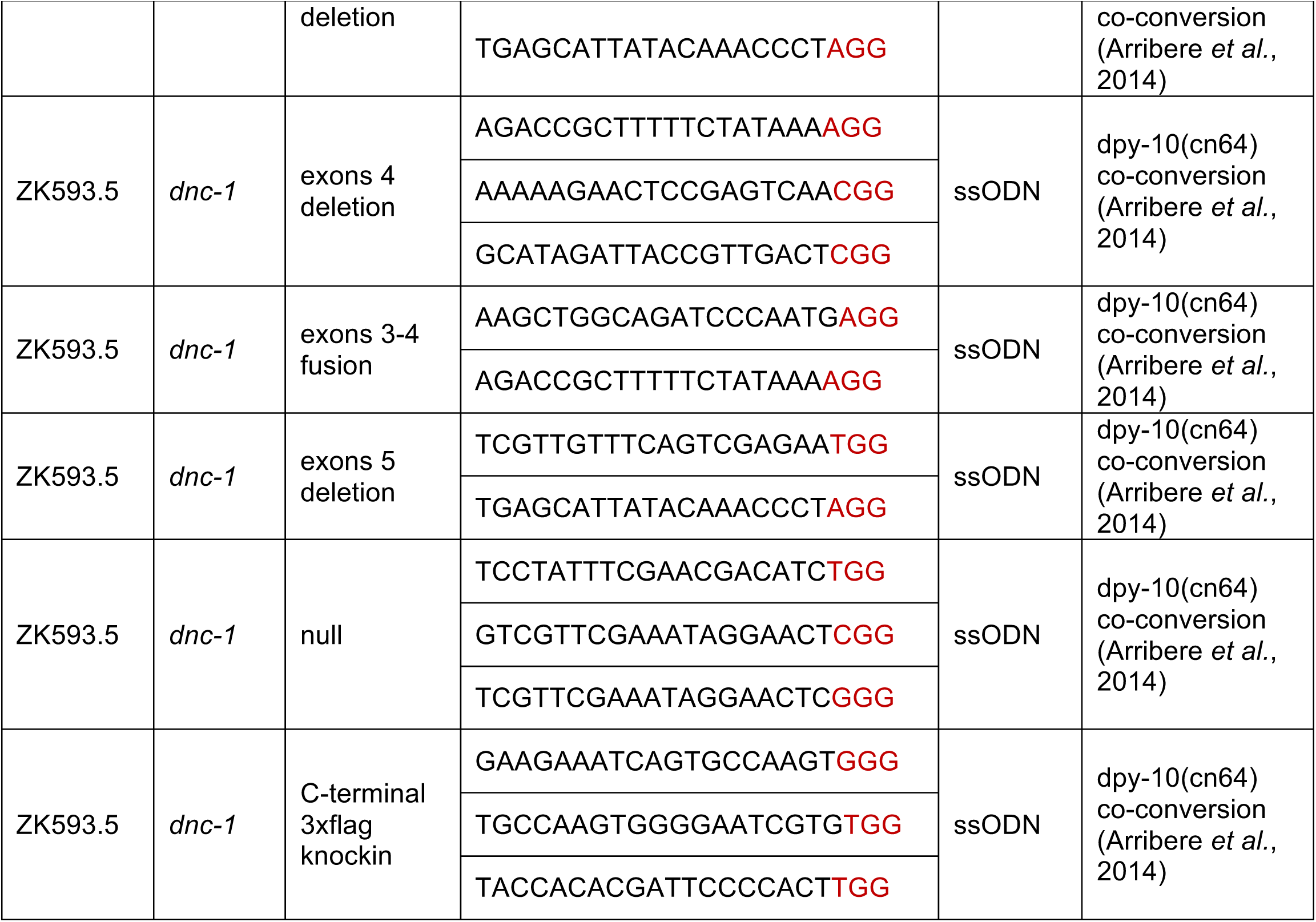
genomic sequences targeted by sgRNAs used for genome editing by CRISPR-Cas9

**Supplemental Table S3:**
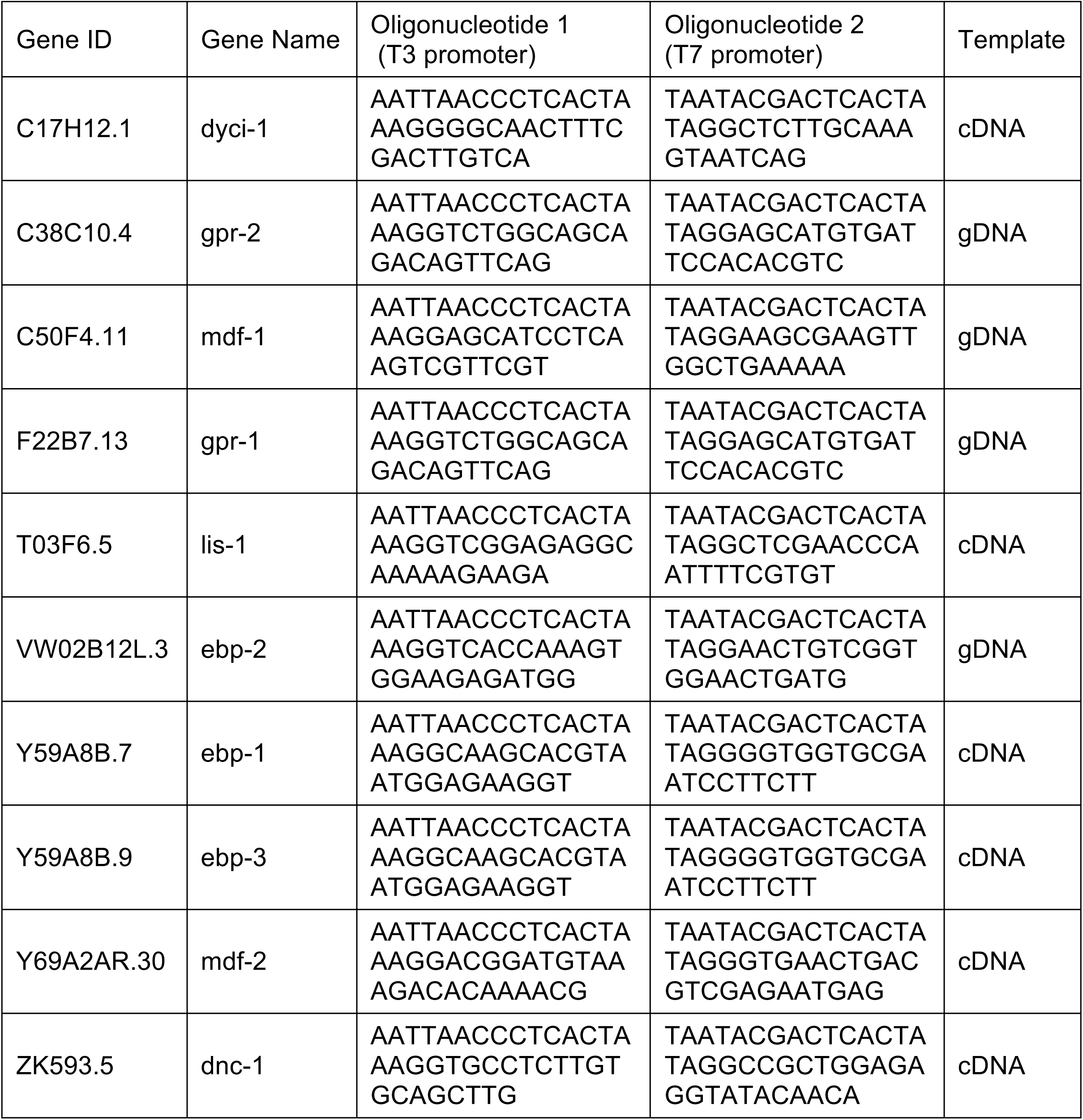
oligos used in this study for double-stranded RNA production

## REFERENCES

Akhmanova A & Steinmetz MO (2015) Control of microtubule organization and dynamics: two ends in the limelight. Nat Rev Mol Cell Biol 16: 711–726

Araki E, Tsuboi Y, Daechsel J, Milnerwood A, Vilariño-Güell C, Fujii N, Mishima T, Oka T, Hara H, Fukae J & Farrer MJ (2014) A Novel DCTN1 mutation with late-onset parkinsonism and frontotemporal atrophy. Mov. Disord.

Arribere JA, Bell RT, Fu BXH, Artiles KL, Hartman PS & Fire AZ (2014) Efficient Marker-Free Recovery of Custom Genetic Modifications with CRISPR/Cas9 in Caenorhabditis elegans. Genetics 198: 837–846

Baugh LR, Hill AA, Slonim DK, Brown EL & Hunter CP (2003) Composition and dynamics of the Caenorhabditis elegans early embryonic transcriptome. Development 130: 889–900

Bieling P, Kandels-Lewis S, Telley IA, van Dijk J, Janke C & Surrey T (2008) CLIP-170 tracks growing microtubule ends by dynamically recognizing composite EB1/tubulin-binding sites. The Journal of Cell Biology 183: 1223–1233

Coffman VC, McDermott MBA, Shtylla B & Dawes AT (2016) Stronger net posterior cortical forces and asymmetric microtubule arrays produce simultaneous centration and rotation of the pronuclear complex in the early Caenorhabditis elegans embryo. Mol Biol Cell 27: 3550–3562

Couwenbergs C, Labbé J-C, Goulding M, Marty T, Bowerman B & Gotta M (2007) Heterotrimeric G protein signaling functions with dynein to promote spindle positioning in C. elegans. J Cell Biol 179: 15–22

Crowder ME, Flynn JR & McNally KP (2015) Dynactin-dependent cortical dynein and spherical spindle shape correlate temporally with meiotic spindle rotation in Caenorhabditis elegans. Molecular biology of …

Culver-Hanlon TL, Lex SA, Stephens AD, Quintyne NJ & King SJ (2006) A microtubule-binding domain in dynactin increases dynein processivity by skating along microtubules. Nat Cell Biol 8: 264–270

De Simone A, Nédélec F & Gönczy P (2016) Dynein Transmits Polarized Actomyosin Cortical Flows to Promote Centrosome Separation. Cell Rep 14: 2250–2262

Desai A, Rybina S, Müller-Reichert T, Shevchenko A, Shevchenko A, Hyman A & Oegema K (2003) KNL-1 directs assembly of the microtubule-binding interface of the kinetochore in C. elegans. Genes & Development 17: 2421–2435

Dixit R, Levy JR, Tokito M, Ligon LA & Holzbaur ELF (2008) Regulation of dynactin through the differential expression of p150Glued isoforms. J Biol Chem 283: 33611– 33619

Duellberg C, Trokter M, Jha R, Sen I, Steinmetz MO & Surrey T (2014) Reconstitution of a hierarchical +TIP interaction network controlling microtubule end tracking of dynein. Nat Cell Biol 16: 804–811

Ellefson MLM & McNally FJF (2011) CDK-1 inhibits meiotic spindle shortening and dynein-dependent spindle rotation in C. elegans. J Cell Biol 193: 1229–1244

Farrer MJ, Hulihan MM, Kachergus JM, Dachsel JC, Stoessl AJ, Grantier LL, Calne S, Calne DB, Lechevalier B, Chapon F, Tsuboi Y, Yamada T, Gutmann L, Elibol B, Bhatia KP, Wider C, Vilariño-Güell C, Ross OA, Brown LA, Castanedes-Casey M, et al (2009) DCTN1 mutations in Perry syndrome. Nat Genet 41: 163–165

Frøkjaer-Jensen C, Davis MW, Ailion M & Jorgensen EM (2012) Improved Mos1-mediated transgenesis in C. elegans. Nat Meth 9: 117–118

Gama JB, Pereira C, Simões PA, Celestino R, Reis RM, Barbosa DJ, Pires HR, Carvalho C, Amorim J, Carvalho AX, Cheerambathur DK & Gassmann R (2017) Molecular mechanism of dynein recruitment to kinetochores by the Rod–Zw10–Zwilch complex and Spindly. J Cell Biol 5: jcb.201610108

Gill SR, Schroer TA, Szilak I, Steuer ER, Sheetz MP & Cleveland DW (1991) Dynactin, a conserved, ubiquitously expressed component of an activator of vesicle motility mediated by cytoplasmic dynein. The Journal of Cell Biology 115: 1639–1650

Gönczy P, Pichler S, Kirkham M & Hyman AA (1999) Cytoplasmic Dynein Is Required for Distinct Aspects of Mtoc Positioning, Including Centrosome Separation, in the One Cell Stage Caenorhabditis elegans Embryo. J Cell Biol 147: 135–150

Hayashi I, Wilde A, Mal TK & Ikura M (2005) Structural basis for the activation of microtubule assembly by the EB1 and p150Glued complex. Mol Cell 19: 449–460

Honnappa S, Okhrimenko O, Jaussi R, Jawhari H, Jelesarov I, Winkler FK & Steinmetz MO (2006) Key interaction modes of dynamic +TIP networks. Mol Cell 23: 663–671

Janke C (2014) The tubulin code: molecular components, readout mechanisms, and functions. The Journal of Cell Biology 206: 461–472

Kim H, Ling S-C, Rogers GC, Kural C, Selvin PR, Rogers SL & Gelfand VI (2007) Microtubule binding by dynactin is required for microtubule organization but not cargo transport. The Journal of Cell Biology 176: 641–651

Kimura A & Onami S (2005) Computer simulations and image processing reveal length-dependent pulling force as the primary mechanism for C. elegans male pronuclear migration. Developmental Cell 8: 765–775

Kimura A & Onami S (2007) Local cortical pulling-force repression switches centrosomal centration and posterior displacement in C. elegans. The Journal of Cell Biology 179: 1347–1354

Kimura K & Kimura A (2011) Intracellular organelles mediate cytoplasmic pulling force for centrosome centration in the Caenorhabditis elegans early embryo. Proceedings of the National Academy of Sciences 108: 137–142

Kotak S & Gönczy P (2013) Mechanisms of spindle positioning: cortical force generators in the limelight. Curr Opin Cell Biol

Kozlowski C, Srayko M & Nédélec F (2007) Cortical microtubule contacts position the spindle in C. elegans embryos. Cell 129: 499–510

Lansbergen G, Komarova Y, Modesti M, Wyman C, Hoogenraad CC, Goodson HV, Lemaitre RP, Drechsel DN, van Munster E, Gadella TWJ, Grosveld F, Galjart N, Borisy GG & Akhmanova A (2004) Conformational changes in CLIP-170 regulate its binding to microtubules and dynactin localization. J Cell Biol 166: 1003–1014

Lenz JH, Schuchardt I, Straube A & Steinberg G (2006) A dynein loading zone for retrograde endosome motility at microtubule plus-ends. EMBO J 25: 2275–2286

Lloyd TE, Machamer J, O’Hara K, Kim JH, Collins SE, Wong MY, Sahin B, Imlach W, Yang Y, Levitan ES, McCabe BD & Kolodkin AL (2012) The p150(Glued) CAP-Gly Domain Regulates Initiation of Retrograde Transport at Synaptic Termini. Neuron 74: 344–360

Markus SM & Lee W-L (2011a) Regulated offloading of cytoplasmic dynein from microtubule plus ends to the cortex. Developmental Cell 20: 639–651

Markus SM & Lee W-L (2011b) Microtubule-dependent path to the cell cortex for cytoplasmic dynein in mitotic spindle orientation. Bioarchitecture 1: 209–215

McKenney RJ, Huynh W, Tanenbaum ME, Bhabha G & Vale RD (2014) Activation of cytoplasmic dynein motility by dynactin-cargo adapter complexes. Science 345: 337– 341

McKenney RJ, Huynh W, Vale RD & Sirajuddin M (2016) Tyrosination of a-tubulin controls the initiation of processive dynein-dynactin motility. EMBO J

Mishima M, Maesaki R, Kasa M, Watanabe T, Fukata M, Kaibuchi K & Hakoshima T (2007) Structural basis for tubulin recognition by cytoplasmic linker protein 170 and its autoinhibition. Proc Natl Acad Sci USA 104: 10346–10351

Moore JK, Sept D & Cooper JA (2009) Neurodegeneration mutations in dynactin impair dynein-dependent nuclear migration. Proceedings of the National Academy of Sciences 106: 5147–5152

Moughamian AJ & Holzbaur ELF (2012) Dynactin is required for transport initiation from the distal axon. Neuron 74: 331–343

Moughamian AJ, Osborn GE, Lazarus JE, Maday S & Holzbaur ELF (2013) Ordered Recruitment of Dynactin to the Microtubule Plus-End is Required for Efficient Initiation of Retrograde Axonal Transport. J. Neurosci. 33: 13190–13203

Nguyen-Ngoc T, Afshar K & Gönczy P (2007) Coupling of cortical dynein and G alpha proteins mediates spindle positioning in Caenorhabditis elegans. Nat Cell Biol 9: 1294–1302

Nirschl JJ, Magiera MM, Lazarus JE, Janke C & Holzbaur ELF (2016) a-Tubulin Tyrosination and CLIP-170 Phosphorylation Regulate the Initiation of Dynein-Driven Transport in Neurons. Cell Rep 14: 2637–2652

Paix A, Wang Y, Smith HE, Lee C-YS, Calidas D, Lu T, Smith J, Schmidt H, Krause MW & Seydoux G (2014) Scalable and Versatile Genome Editing Using Linear DNAs with Micro-Homology to Cas9 Sites in Caenorhabditis elegans. Genetics

Pecreaux J, Röper J-C, Kruse K, Jülicher F, Hyman AA, Grill SW & Howard J (2006) Spindle Oscillations during Asymmetric Cell Division Require a Threshold Number of Active Cortical Force Generators. Current Biology 16: 2111–2122

Peris L, Thery M, Fauré J, Saoudi Y, Lafanechère L, Chilton JK, Gordon-Weeks P, Galjart N, Bornens M, Wordeman L, Wehland J, Andrieux A & Job D (2006) Tubulin tyrosination is a major factor affecting the recruitment of CAP-Gly proteins at microtubule plus ends. J Cell Biol 174: 839–849

Puls I, Jonnakuty C, LaMonte BH, Holzbaur ELF, Tokito M, Mann E, Floeter MK, Bidus K, Drayna D, Oh SJ, Brown RH, Ludlow CL & Fischbeck KH (2003) Mutant dynactin in motor neuron disease. Nat Genet 33: 455–456

Reinsch S & Gönczy P (1998) Mechanisms of nuclear positioning. Journal of Cell Science 111: 2283–2295

Rose L & Gönczy P (2014) Polarity establishment, asymmetric division and segregation of fate determinants in early C. elegans embryos. WormBook: 1–43

Schlager MA, Hoang HT, Urnavicius L, Bullock SL & Carter AP (2014) In vitro reconstitution of a highly processive recombinant human dynein complex. EMBO J 33: 1855–1868

Schmidt DJ, Rose DJ, Saxton WM & Strome S (2005) Functional analysis of cytoplasmic dynein heavy chain in Caenorhabditis elegans with fast-acting temperature-sensitive mutations. Mol Biol Cell 16: 1200–1212

Schroer TA (2004) Dynactin. Annu. Rev. Cell Dev. Biol. 20: 759–779

Schroer TA & Sheetz MP (1991) Two activators of microtubule-based vesicle transport. The Journal of Cell Biology 115: 1309–1318

Shinar T, Mana M, Piano F & Shelley MJ (2011) A model of cytoplasmically driven microtubule-based motion in the single-celled Caenorhabditis elegans embryo. Proceedings of the National Academy of Sciences 108: 10508–10513

Splinter D, Razafsky DS, Schlager MA, Serra-Marques A, Grigoriev I, Demmers J, Keijzer N, Jiang K, Poser I, Hyman AA, Hoogenraad CC, King SJ & Akhmanova A (2012) BICD2, dynactin, and LIS1 cooperate in regulating dynein recruitment to cellular structures. Mol Biol Cell 23: 4226–4241

Steinmetz MO & Akhmanova A (2008) Capturing protein tails by CAP-Gly domains. Trends Biochem Sci 33: 535–545

Tanimoto H, Kimura A & Minc N (2016) Shape–motion relationships of centering microtubule asters. J Cell Biol 212: 777–787

Tokito MK, Howland DS, Lee VM & Holzbaur EL (1996) Functionally distinct isoforms of dynactin are expressed in human neurons. Mol Biol Cell 7: 1167–1180

Vaughan KT, Tynan SH, Faulkner NE, Echeverri CJ & Vallee RB (1999) Colocalization of cytoplasmic dynein with dynactin and CLIP-170 at microtubule distal ends. Journal of Cell Science 112 (Pt 10): 1437–1447

Vaughan PS, Miura P, Henderson M, Byrne B & Vaughan KT (2002) A role for regulated binding of p150(Glued) to microtubule plus ends in organelle transport. The Journal of Cell Biology 158: 305–319

Walston TD & Hardin J (2006) Wnt-dependent spindle polarization in the early C. elegans embryo. Seminars in cell & developmental biology 17: 204–213

Wang Q, Crevenna AH & Kunze I (2014) Structural basis for the extended CAP-Gly domains of p150glued binding to microtubules and the implication for tubulin dynamics. In

Waterman-Storer CM, Karki S & Holzbaur EL (1995) The p150Glued component of the dynactin complex binds to both microtubules and the actin-related protein centractin (Arp-1). Proc Natl Acad Sci USA 92: 1634–1638

Watson P & Stephens DJ (2006) Microtubule plus-end loading of p150Glued is mediated by EB1 and CLIP-170 but is not required for intracellular membrane traffic in mammalian cells. Journal of Cell Science 119: 2758–2767

Weisbrich A, Honnappa S, Jaussi R, Okhrimenko O, Frey D, Jelesarov I, Akhmanova A & Steinmetz MO (2007) Structure-function relationship of CAP-Gly domains. Nat Struct Mol Biol 14: 959–967

Wühr M, Tan ES, Parker SK, Detrich HW & Mitchison TJ (2010) A model for cleavage plane determination in early amphibian and fish embryos. Curr Biol 20: 2040–2045

Yao X, Zhang J, Zhou H, Wang E & Xiang X (2012) In vivo roles of the basic domain of dynactin p150 in microtubule plus-end tracking and dynein function. Traffic 13: 375– 387

Yu I, Garnham CP & Roll-Mecak A (2015) Writing and Reading the Tubulin Code. Journal of Biological Chemistry 290: 17163–17172

Zhang H, Skop AR & White JG (2008) Src and Wnt signaling regulate dynactin accumulation to the P2-EMS cell border in C. elegans embryos. Journal of Cell Science 121: 155–161

Zhang J, Zhuang L, Lee Y, Abenza JF, Peñalva MA & Xiang X (2010) The microtubule plus-end localization of Aspergillus dynein is important for dynein-early-endosome interaction but not for dynein ATPase activation. Journal of Cell Science 123: 3596– 3604

Zhapparova ON, Bryantseva SA, Dergunova LV, Raevskaya NM, Burakov AV, Bantysh OB, Shanina NA & Nadezhdina ES (2009) Dynactin subunit p150Glued isoforms notable for differential interaction with microtubules. Traffic 10: 1635–1646

